# Transformation of primary murine peritoneal mast cells by constitutive KIT activation as a result of lost *Cdkn2a/Arf* expression

**DOI:** 10.1101/2023.01.24.525344

**Authors:** Sandro Capellmann, Roland Sonntag, Herdit Schüler, Steffen K Meurer, Lin Gan, Marlies Kauffmann, Katharina Horn, Hiltrud Königs-Werner, Ralf Weiskirchen, Christian Liedtke, Michael Huber

## Abstract

Mast cells (MCs) are immune cells of the myeloid lineage distributed in tissues throughout the body. Phenotypically, they are a heterogeneous group characterized by different protease repertoires stored in secretory granules and differential presence of receptors. To adequately address aspects of MC biology either primary MCs isolated from human or mouse tissue or different human MC lines, like HMC-1.1 and -1.2, or rodent MC lines like L138.8A or RBL-2H3 are frequently used. Nevertheless, cellular systems to study MC functions are very limited. We have generated a murine connective tissue-like MC line, termed PMC-306, derived from primary peritoneal MCs (PMCs), which spontaneously transformed. We analyzed PMC-306 cells regarding MC surface receptor expression, effector functions and respective signaling pathways, and found that the cells reacted very similar to primary wildtype (WT) PMCs. In this regard, stimulation with MAS-related G-protein-coupled receptor member B2 (MRGPRB2) ligands induced respective signaling and effector functions. Furthermore, PMC-306 cells revealed significantly accelerated cell cycle progression, which however was still dependent on IL-3 and stem cell factor (SCF). Phenotypically, PMC-306 cells adopted an immature connective tissue-like MCs appearance. The reason for immortalization most likely is the loss of the two critical cell cycle regulators *Cdkn2a*/INK4A and *Arf*/p19, respectively. The loss of *Cdkn2a* and *Arf* expression could be mimicked in primary bone marrow-derived mast cells (BMMCs) by SCF supplementation strongly arguing for an involvement of KIT activation in the transformation process. Hence, this new cell line might be a useful tool to study further aspects of PMC function and to address tumorigenic processes associated with MC leukemia.

## Introduction

Mast cells (MCs) are tissue-resident immune cells of myeloid origin and well-known effectors of type I hypersensitivity (1–3). Besides, MCs can contribute to cancer progression by supporting angiogenesis, extracellular matrix degradation, and epithelial-to-mesenchymal transition (4, 5). The immune response to viral and bacterial infections is fundamentally triggered by MCs as they are strategically located at interfaces between host and environment (3, 6). Therefore, they belong to the first line of defence against intruders and fulfil the task to alert other immune cells to migrate to infected tissues (7, 8). Finally, MCs also contribute to tissue repair and homeostasis in the resolution phase of inflammation by secreting anti-inflammatory IL-10 and helping to remodel the extracellular matrix (9–11).

Understanding fundamental MC functions is often impeded by adequate cellular model systems. One reason is that a prototypical MC does not exist. The term mast cell comprises a heterogeneous cell population that can be classified into multiple subgroups depending on the receptor repertoire, composition of secretory granules and tissue residency (12). Thus, finding *in vitro* models that represent all aspects of different MC subtypes is not possible. Different tissue sources used for isolation and differences concerning *in vitro* differentiation and/or culturing of MCs using various cytokine and growth factor combinations influences analyses of general MC features. Due to the difficulties to isolate mature tissue-derived MCs from human or murine sources, many studies dealing with MCs have been conducted using human MC lines like HMC-1, LUVA and LAD2 (13–15) or rodent MC lines like RBL-2H3 or L138.8A (16, 17). However, results obtained with these cell lines have to be interpreted with caution. Immortalized leukemic cell lines, like the HMC-1 line usually contain oncogenic mutations, e.g. in the *KIT* gene (18), which influences studies on MC biology. Furthermore, the HMC-1 cell line does not express a functional FcεR1 on the cell surface impeding studies on IgE receptor signaling. Likewise, the RBL-2H3 cell line, for instance, shows substantial genomic rearrangements whose consequences are scarcely characterized but certainly affect signaling pathways and cellular physiology in multiple ways (19, 20).

In mouse and rats, MCs are – in a simplified way – divided into two groups based on their histochemical staining properties reflecting different granule mediator content. Connective tissue MCs (CTMCs) typically contain tryptase, chymases, carboxypeptidase A3 (CPA3) and heparin proteogylcans, whereas granules of mucosal MCs (MMCs) lack tryptase and instead of heparin predominantly contain chondroitin sulfate (21). *In vitro*, murine mucosal-like MCs can be obtained from multipotent bone marrow-derived precursor cells, which require a long-lasting differentiation process in the presence of IL-3 and (optionally) stem cell factor (SCF). The resulting MCs are referred to as bone marrow-derived MCs (BMMCs). However, *in vitro* differentiated cultures of murine BMMC frequently result in an incompletely differentiated population of MCs, which morphologically and histochemically do not fully correspond to finally differentiated MCs observed in mouse tissues (22). Connective tissue-like MCs can be isolated from peritoneum – hence they are referred to as murine peritoneum-derived MCs (PMCs) – and subsequently enriched and expanded in culture in the presence of IL-3 and SCF (23). Challenging about PMCs is, however, that peritoneal lavages often yield only strictly limited amounts of enriched PMCs that cannot be cultured over a longer time period. As a consequence, to extensively characterize PMC biology, plenty of mice need to be sacrificed to yield validated scientific data, which is not in agreement with the 3R principles of animal welfare (24).

To obtain a comprehensive picture of general MC characteristics and functions, a combined approach analyzing cell lines, primary cells and *in vivo* models is required. In the present study, we describe the generation of a PMC-derived spontaneously transformed murine MC line termed PMC-306 that we propose as a new additional tool to study MC biology. We analyzed the MC characteristics in terms of morphology, cell surface marker expression, pro-inflammatory effector responses to different stimuli and protease expression in relation to wildtype (WT) PMCs. We could reveal that the PMC line retained its MC features. PMC-306 cells express functional high-affinity IgE and MRGPRB2 receptors and can therefore be used for respective signaling studies. In addition, we investigated proliferation, survival and the cell cycle pattern of PMC-306, which showed dependence on the cytokines SCF and IL-3. The chromosomal analysis did not reveal major chromosomal abnormalities apart from a heterozygous deletion of regions qC4-qC7 on chromosome 4. This region, amongst others, encodes the cyclin-dependent kinase inhibitor 2A (*Cdkn2a*, also known as INK4A or p16) and *Arf* (also known as p19), both of which function as central cell cycle inhibitors regulating the p53 and Rb pathways (25, 26). Intriguingly, despite the observed heterozygosity of the change on DNA level, mRNA expression of *Cdkn2a* and *Arf* was not detectable in PMC-306 cells. We demonstrated that these aberrations lead to an enhanced cell cycle activity, a strongly increased proliferative capacity and best explains the mechanism of PMC transformation. We could further induce loss of *Cdkn2a* and *Arf* expression by SCF supplementation of BMMC cultures pointing to a role of chronic KIT activation in the immortalization process. Our initial characterization of PMC-306 provides the basis for using this cell line in prospective studies as a new tool to address differential MC functions.

## Material and Methods

### Animals and Cell Culture

Murine peritoneal MCs (PMCs) and bone marrow-derived MCs (BMMCs) were isolated and cultured as described previously (23). Primary WT PMCs were cultured in conventional cell culture medium (RPMI1640 (Invitrogen, #21875-0991) containing 15% FCS (Capricorn, #FBS-12A), 10 mM HEPES (pH 7.4), 50 units/ml Penicillin and 50 mg/ml Streptomycin, 100 µM β-mercaptoethanol, 30 ng/ml IL-3 from X63-Ag8-653 conditioned medium (27) and approximately, depending on its biological activity, 20 ng/ml SCF from CHO culture supernatant (23)). For cultivation of PMC-306 cells, the amounts of FCS and SCF were reduced from 15% to 10% and from 20 ng/ml to 5 ng/ml, respectively. Primary BMMCs were cultivated like PMCs, however, SCF was only supplemented to analyze the influence of KIT activation. PMC-306 cells were frozen at -150°C in 90% FCS and 10% DMSO in aliquots of 10 x 10^6^ cells per ml. WT PMCs were freshly isolated and were cultured for a maximum of 8 weeks. All mice used in this study were on a mixed C57BL/6 x 129/Sv background. Experiments were performed in accordance with German legislation governing animal studies and following the principles of laboratory animal care. Mice were held in the Institute of Laboratory Animal Science, Medical Faculty of RWTH Aachen University. The institute holds a license for husbandry and breeding of laboratory animals from the veterinary office of the Städteregion Aachen (administrative district). The institute follows a quality management system, which is certified according to DIN EN ISO 9001:2015. Every step in this project involving mice was reviewed by the animal welfare officer.

### **β**-Hexosaminidase Assay

Measuring FcεRI-dependent degranulation, MCs were preloaded with 0.15 µg/ml IgE (clone Spe-7, Sigma, #A2831) overnight (37°C, 5% CO_2_). This step was skipped when cells were stimulated *via* MRGPRB2. Cells were washed in sterile PBS, resuspended at a density of 1.2 x 10^6^ (PMC-306 line) or 0.6 x 10^6^ (WT PMCs) per ml in Tyrode’s buffer (130 mM NaCl, 5 mM KCl, 1.4 mM CaCl_2_, 1 mM MgCl_2_, 5.6 mM glucose, and 0.1% BSA in 10 mM HEPES, pH 7.4) and allowed to adapt to 37°C 15 min. For stimulation, different concentrations of antigen (Ag; DNP-HSA, Sigma, #A6661), Mastoparan (Sigma, #M5280) or Compound 48/80 (C48/80) (Sigma, #C2313) were applied for 30 minutes. The amount of released β-hexosaminidase was determined as described in (28).

### Cytokine ELISA

To determine IL-6 and TNF secretion, WT PMCs and PMC-306 were stimulated as indicated in the respective experiments. If cells were stimulated with Ag, MCs were pre-loaded with IgE. Cell number was adjusted to 1.2 x 10^6^/ml in stimulation medium (RPMI 1640 + 0.1% BSA (Serva, #11930.04)), cells were allowed to adapt to 37°C and stimulated for 3 hours. Supernatants were collected after centrifugation. 96-well ELISA plates (Corning, #9018) were coated with capturing anti-IL-6 (1:250, BD Biosciences, #554400) or anti-TNF (1:200, R&D Systems, #AF410-NA) antibodies diluted in PBS overnight at 4°C. ELISA plates were washed three times with PBS+0.1% Tween and subsequently blocked with PBS + 2% BSA (IL-6 ELISA) or PBS + 1% BSA + 5% sucrose (TNF ELISA) for 90 min before loading the supernatants (50 µl for IL-6 ELISA, 100 µl for TNF ELISA). Additionally to loading of supernatants, IL-6 (BD Pharmingen, #554582) and TNF (R&D Systems, #410-MT-010) standards in 1:2 dilutions were added and plates were incubated overnight at 4°C. Thereupon, plates were washed three times again followed by incubation with biotinylated anti-IL-6 (1:500, BD Biosciences, #554402) and anti-TNF antibodies (1:250, R&D Systems, #BAF-410) diluted in PBS+1% BSA for 45 minutes and 2 hours, respectively, at room temperature (RT). After 3 washing steps, Streptavidin alkaline phosphatase (SAP, 1:1000, BD Pharmingen, #554065) was added for 30 minutes at RT. After 3 more washing steps, the substrate p-Nitro-phenyl-phosphate (1 pill per 5 ml in sodium carbonate buffer (2 mM MgCl_2_ in 50 mM sodium carbonate, pH 9.8), Sigma, #S0942-200TAB) was added and OD_450_ was recorded using a plate reader (BioTek Eon).

### Flow Cytometry, Viability Assay, LAMP-1 Assay and Cell Cycle Analysis

Flow cytometry was performed to analyze MC surface marker expression. Roughly 0.5 x 10^6^ cells were washed in FACS buffer (PBS + 3% FCS + 0.1% sodium azide) and stained with either FITC-coupled anti-FcεR1 (1:100, clone MAR-1, BioLegend, #134306) and PE-coupled anti-CD117 (1:100, clone 2B8, BioLegend, #105808) or FITC-coupled anti-ST2 (1:100, clone DJ8, MD Bioproducts, #101001F) and PE-coupled anti-CD13 (1:100, clone R3-242, BD Pharmingen, #558745), respectively, for 20 minutes at 4°C in the dark. Thereupon, cells were washed again in FACS buffer, resuspended in 200 µl FACS buffer and analyzed using a FACSCanto II (BD Biosciences).

To determine viability of WT PMCs and PMC-306 cells, cells were seeded at a density of 0.3 x 10^6^ (PMC-306) or 0.5 x 10^6^ (WT PMCs) cells/ml and incubated under different cytokine deprivation conditions for 72 hours. Thereupon, cells were washed in FACS buffer and stained with Annexin V (1:100, BioLegend, #640912) diluted in culture medium for 20 minutes at 4°C in the dark. Immediately before analysis by flow cytometry using the FACSCanto II, propidium iodide (Sigma, #P4864) at a concentration of 1 µg/ml was added.

For LAMP-1 assay, MCs were preloaded with IgE (Spe7) overnight. Cells were washed and cell number was adjusted to 1 x 10^6^ cells per ml in RPMI 1640 + 0.1% BSA. Stimulation was performed for indicated time points with Ag (DNP-HSA). The reaction was stopped by centrifugation at 4°C followed by staining of cell surface LAMP-1 (CD107a) using FITC-coupled anti-LAMP-1 antibody (clone 1D4B, BioLegend, #121605) for 20 minutes at 4°C in the dark. Cells were washed in FACS buffer and LAMP-1 cell surface expression was determined by flow cytometry using a FACSCanto II.

For analysis of cell cycle parameters, background values of pooled (from n=3 per PMC cell type), unstained cells were subtracted from values of three independent stained samples. Cultured cells were pelleted by centrifugation (1400 rpm, 4 °C, 10 min), subsequently stained with fluorophore-labelled antibodies against the surface markers CD117-PE-Cy7 (1:100, # 561681, BD Biosciences) and FcεR1-FITC (1:100, #11-5895-81, Invitrogen Thermo Fisher Scientific) in FACS-blocking buffer (mixture of 0.66% human/rabbit/mouse serum, Sigma-Aldrich, and 1% Bovine Serum Albumin, Sigma-Aldrich) for 40 minutes at 4 °C. Cell fixation and permeabilization was performed using the eBioscience Foxp3/Transcription Factor Staining Buffer Set (Invitrogen Thermo Fisher Scientific) according to the manufacturer’s protocol. Intracellular co-staining of cells was performed using a mix of fluorophore-labelled antibodies against pH2Ax-PerCP-Cy5.5 (1:100, #564718, S139, BD Biosciences) together with pH3-AL647 (1:100, #3458, S10, Cell Signaling Technology) or MKi67-AL700 (1:100, #56-5698-82, Invitrogen Thermo Fisher Scientific) diluted in FACS-blocking buffer (mixture of 0.66% human/rabbit/mouse serum, Sigma-Aldrich, and 1% Bovine Serum Albumin, Sigma-Aldrich in PBS) for 30 minutes at 4°C. The cellular DNA content was determined by DAPI staining (BD Pharmingen, 1:1000 in 1x PBS). Compensation of each fluorochrome was automatically performed using OneComp ebeads (#01-111-42, Invitrogen Thermo Fisher Scientific) according to manufacturer’s recommendations. Measurements were performed using a BD LSRFortessa (BD Biosciences).

Acquired flow cytometry data were analyzed with FlowJo software v10 (Tree Star).

### RT-qPCR

Total RNA was isolated from 4 x 10^6^ cells using NucleoSpin RNA Plus Kit (Macherey Nagel, #740955.50) according to manufacturer’s instructions. 1 µg of RNA was reverse transcribed using random oligonucleotides (Roche, #11034731.001) and Omniscript Reverse Transcription (RT) Kit (Qiagen, #205113) according to manufacturer’s instructions. Quantification of transcript expression was performed using Sybr green reaction mix SensiFAST (Bioline, #BIO-86005) and 10 pmol of specific primers. PCR reaction was performed on a Rotorgene Q (Qiagen). Transcript expression was normalized to the housekeeping gene *Hprt* and relative expression was calculated according to the delta-C_T_ method (29). Following primers were used: *Hprt-*fwd: GCTGGTGAAAAGGACCTCC, *Hprt-*rev: CACAGGACTAGAACACCTGC, *Cdkn2a-*fwd: CTTTCGGTCGTACCCCGATT, *Cdkn2a-*rev: AGAAGGTAGTGGGGTCCTCG, *Arf-*fwd: TGGTGAAGTTCGTGCGATCC, *Arf-*rev: TACGTGAACGTTGCCCATCA, *Rbl1-*fwd: AACTGAACCTGGACGAGGGA, *Rbl1-*rev: GAGCATGC CAGCCAGTGTAT, *Dennd4c-*fwd: GGGAGAGACTCTGTCGCCTA, *Dennd4c-*rev: AACGTCTCCACTGCTGCTAC; *Mllt3-*fwd: ATGGCTAGCTCGTGTGCC, *Mllt3-*rev: GAACACCATCCAGTCGTGGG, *Plaa-*fwd: CAGACAGTCCTAACAGGGGC, *Plaa-* rev: TCCTCCAGTGGCAATCAGTC, *Cpa3-*fwd: AATTGCTCCTGTCCACTTTGA, *Cpa3-*rev: TCACTAACTCGGAAATCC ACAGT, *Gzmb-*fwd: ATGGCCCCAATGGGCAAATA, *Gzmb-*rev: CCGAAAGGAAGCACGTTTGG, *Mcpt2-* fwd: TTCACCACTAAGAACG GTTCG, *Mcpt2-*rev: CTCCAAGGATGACACTGATTTCA, *Mcpt4-*fwd: GTGGGC AGTCCCAGAAAGAA, *Mcpt4-*rev: GCATCTCCGCGTCCATAAGA, *Tpsab1 (Mcpt7)-* fwd: GCCAATGACACCTACTGGATG, *Tpsab1 (Mcpt7)-*rev: GAGCTGTACT CTGACCTTGTTG, *Cma1-*fwd: ACGGACAGAGGTTCTGAGGA, *Cma1-*rev: GAGCTCCAAGGGTGACAGTG, *Mrgprb2-*fwd: CCTCAGCCTGGAAAACGAAC, *Mrgprb2-*rev: CCATCCCAACCAGGGAAATGA, *Ccnd1-*fwd: TCAAGACG GAGGAGACCTGT, *Ccnd1-*rev: GGAAGCGGTCCAGG TAGTTC.

### Stimulation of MCs, Western Blotting and Antibodies

For Ag-dependent MC stimulation, cells were preloaded with IgE (0.15 µg/ml) overnight. For FcεRI-independent stimulation, cells were not preloaded with IgE. Cells were washed with sterile PBS and concentration was adjusted to 1 x 10^6^ cells per ml in RPMI1640 + 0.1% BSA. 1 x 10^6^ cells were stimulated as indicated. Stimulation was stopped by snap-freezing in liquid nitrogen and subsequent lysis in phosphorylation solubilization buffer (PSB) (50 mM HEPES, 100 mM sodium fluoride, 10 mM sodium pyrophosphate, 2 mM sodium orthovanadate, 2 mM EDTA, 2 mM sodium molybdate, 0.5% NP-40, 0.5% sodium deoxycholate and 0.03% sodium dodecylsulfate (SDS)) for 30 minutes at 4°C. Cell lysates were centrifuged (10 min, 13,0000 x g) and subjected to SDS-PAGE and Western blot analysis as described previously (30). The following antibodies were used for detection of p-PLCγ1 (Y783, Cell Signaling Technology (CST), #2821), PLCγ1 (CST, #2822), p-PKB (S473, CST, #9271), PKB (CST, #9272), p-ERK1/2 (T202/Y204, CST, #4370), ERK2 (Santa Cruz, #sc-1647), ERK1/2 (CST, #4696), p-p38 (T180/Y182, CST, #9216), p38 (Santa Cruz, #sc-81621), p-IκBα (S32, CST, #2859), IκBα (CST, #4812), p-KIT (Y719, CST, #3391), KIT (CST, #3074), GAPDH (Santa Cruz, #166574), HSP 90 (CST, #4877), actin (Santa Cruz, #sc-8432), granzyme B (CST, #4275) and tryptase (Santa Cruz, #sc-32889). Secondary antibodies coupled to horseradish peroxidase were purchased from Dako Cytomation (goat anti-rabbit HRP (#P0448), rabbit anti-mouse HRP (#P0161)).

### Proliferation and XTT Assay

Cells were seeded at a density of 0.3 x 10^6^ cells per ml (PMC-306) or 0.5 x 10^6^ cells per ml (WT PMCs) in medium containing different amounts of cytokines and growth factors or different amounts of Imatinib (Selleckchem, #S1026) using DMSO (Applichem, #A3072) as control. After 24 hours of incubation in a humidified atmosphere (37°C, 5% CO_2_), cells were resuspended and 50 µl of cell suspension were diluted in 10 ml PBS. Cell count and viability was determined using Casy cell counter (Innovatis). Cell count was monitored over a period of 72 hours.

Metabolic activity was determined using XTT Cell proliferation kit II (Roche, #11465015001). Cells were seeded at a density of 0.3 x 10^6^ (PMC-306) or 0.5 x 10^6^ (WT PMCs) cell per ml in wells of a 96-well microplate and a final volume of 100 µl. Metabolic activity was measured after an incubation time of 72 hours under humidified culture conditions (37°C, 5% CO_2_) by adding 50 µl of XTT reagent to each well and measuring spectrophotometrical absorbance of the resulting formazan product at a wavelength of 475 nm and a reference wavelength of 650 nm. For analysis, the measured absorbances at 475 nm and 650 nm were blanked to medium controls and for total absorbance the blanked absorbance at 475 nm was subtracted from blanked absorbance at 650 nm (blanked A_475_ – blanked A_650_). These calculated values are provided in the figures.

### Electron Microscopy

5 x 10^6^ primary WT PMCs or transformed PMC-306 were centrifuged and resuspended in 500 µl regular growth medium. For fixation, 3% glutaraldehyde dissolved in PBS was added for four hours. The subsequent steps for sample preparations and image acquisition were conducted by the Department of Electron Microscopy at the Institute of Pathology at the University Hospital Aachen. Briefly, cells were centrifuged (2000 rpm, 5 min) and washed in 0.1 M Soerensen’s phosphate buffer (0.133 M Na_2_HPO_4_, 0.133 M KH_2_PO_4_). Post-fixation was performed in 1% OsO_4_. After washing the cells in deionized water, the samples were dehydrated by an ethanol series (30%, 50%, 70%, 90%, and 100%) for 10 minutes each. The specimens were then embedded in agarose (0.25 g in 10 ml deionized water) by stirring the fixed cells into liquid agarose at 60°C. Agarose was allowed to polymerize completely at room temperature and finally cut into multiple blocks that were then embedded in pure Epon resin at 37°C for 12 hours and at 80°C for 48 hours. Specimens were analyzed using a Zeiss Leo906 electron microscope operating at 60 kV.

### May-Grünwald-Giemsa (MGG) Staining

1 x 10^6^ primary PMCs were centrifuged (1,200 rpm, 5 min, 21°C) and resuspended in 100 µL PBS. 50 µL of the cell suspension was spotted onto an ethanol cleaned glass slide (microscope slides, 76 x 26 mm, pre-cleaned, R. Langenbrink, Emmendingen, Germany) and distributed equally. The glass slide was thoroughly air dried. For the cell line PMC-306 1.5 x 10^6^ cells were centrifuged (800 rpm, 5 min, 21°C) and resuspended in 120 µL PBS. 40 µL of the cell suspension was spotted onto an ethanol cleaned glass slide and distributed equally. The glass slide was thoroughly air dried. The cells were fixed by dipping 10 times in pre-cooled ice-cold acetone (VWR, #20066.296) in a glass tray. Thereafter, the glass slide was thoroughly air dried. The staining of MCs with the MGG stain was performed automated according to standard procedures. Microscopic analysis was done by using a slide scanner device (“Fritz” slide scanner, PreciPoint, Germany) equipped with a 40 x objective and the software MicroPoint (V.2021-01). Higher resolution was obtained by digital zoom. To view or transfer data, the software ViewPoint (V1.0.0.9628) and ConvertPoint (V1.0.0.299) were used.

### Cytogenetic Analysis

Structural and numerical chromosome alterations were investigated by conventional karyotyping of the PMC-306 cell line using GTG banding at a 400 to 550 band level (31). Standard procedure to obtain metaphase spreads were performed (32). Cell division was blocked at metaphase followed by hypotonic treatment and methanol/acetic acid fixation (3:1). The banding techniques included use of a trypsin pre-treatment, which was done according to standard protocols (33). Microscopic evaluation was performed using Axioplan fluorescence microscope (Carl Zeiss) and IKARUS digital imaging systems (MetaSystems). 18 to 24 GTG banded metaphases were analyzed.

### Next Generation Sequencing

5 x 10^6^ cells were harvested directly from the respective cell cultures of primary WT PMCs and PMC-306 and washed in PBS. Cell pellets were resuspended in RNAlater buffer (Qiagen, #1017980) and stored at -80°C. The quantity of RNA was analyzed with the Quantus Fluorometer (Promega, Madison, USA). RNA quality control was done using the 4200 TapeStation System (Agilent Technologies, Inc., Santa Clara, USA). An RNA Integrity Number (RIN) of at least 7.3 verified the high quality of all included RNA samples. Sequencing libraries were generated from 1000 ng of total RNA using TruSeq Stranded Total RNA Library Preparation Kit (Illumina, San Diego, USA, #20020596). The libraries were run on an Illumina NextSeq500 High Output 150 cycles Kit v2.5 (Illumina, San Diego, USA, #20024906). FASTQ files were generated using bcl2fastq (Illumina) with standard parameters.

### Statistical Analysis

All data shown were generated from at least three independent experiments. Statistical analysis and graphing of data was performed using GraphPad Prism 9 (GraphPad Software, San Diego, CA 92108). All statistical test procedures were done as described in the respective figure legends. *P*-values were considered statistically significant according to the GP style in GraphPad Prism (ns: *p*>0.05, * *p*<0.05, ** *p*<0.01, *** *p*<0.001, **** *p*<0.0001). The respective number of independent biological replicates per experiment is indicated in the figure legends.

## Results

The murine peritoneal MC line PMC-306 spontaneously developed from murine WT peritoneum-derived MCs. Primary PMCs were isolated and enriched from mice with a mixed C57BL/6;129Sv genetic background as described previously (23). After a culture period of 3 weeks, purity of the parental PMC culture was verified. The homogenous cell population showed ≥ 90% double positivity for FcεRI and KIT/CD117 (suppl. Fig. 1A). While usually after a maximum of 8 weeks in culture, PMC proliferation and viability decreases, PMC-306 cells showed accelerated proliferation with increasing passages. Besides, PMC-306 could be cryopreserved for storage and thawed to be taken in culture again, which is not possible with differentiated primary PMCs or BMMCs. These observations led us to characterize the phenotype of the PMC-306 cell line.

**Fig. 1:**
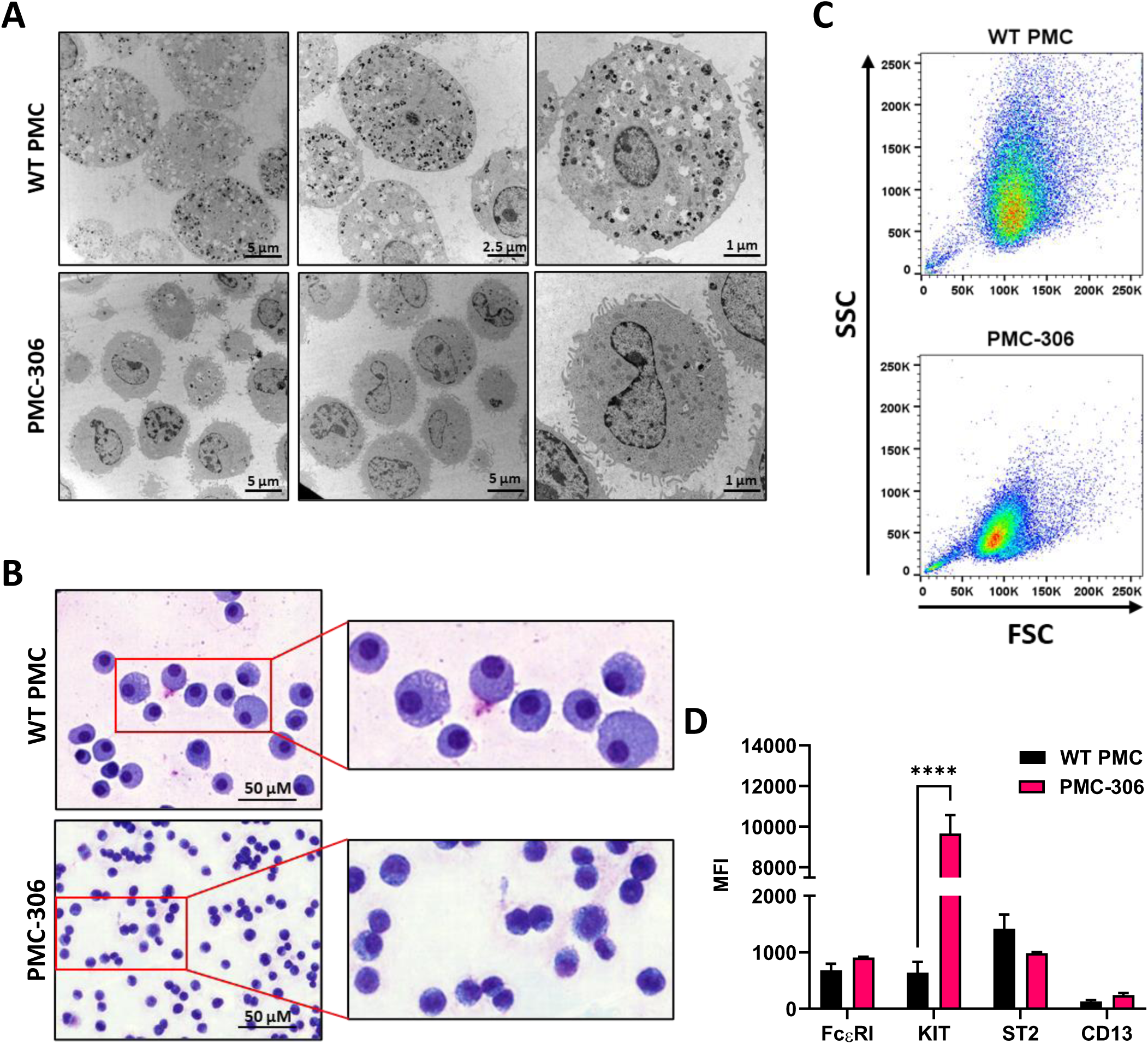
Phenotypic characterization of PMC-306 cells reveals morphological differences to primary WT PMCs. **(A)** Representative electron micrographs of glutaraldehyde-fixed and agarose embedded primary WT PMCs and PMC-306 cells. **(B)** Representative microscopy images of fixed MGG-stained primary WT PMCs and PMC-306 cells. **(C)** FACS analysis of size and granularity of primary WT PMCs in comparison to PMC-306. Forward/side scatter intensities of 30,000 cells were recorded with the same settings to show differences between the two cell types. **(D)** FACS analysis of the MC surface markers FcεRI, KIT (CD117), ST2 and CD13. Geometric mean fluorescence intensities (MFI) are plotted for each surface marker (n=4). Data are shown as mean + SD. Ordinary two-way ANOVA followed by Sídák multiple comparisons test. **** *p*<0.0001.

### PMC-306 cells are smaller and less granulated than primary WT PMCs

One hallmark of MCs are their secretory granules containing, amongst others, biogenic amines and proteases, which can be released upon stimulation in the process of degranulation (34). Electron microscopy analysis revealed drastically reduced granularity of PMC-306 cells compared to primary PMCs, which showed a cytoplasm densely packed with electron dense and seemingly empty granular structures (Fig. 1A). Besides, the nucleus of PMC-306 was enlarged compared to primary PMCs. Similarly, primary PMCs exhibited an increased cytoplasm-to-nucleus relation and were larger compared to PMC-306 as shown by MGG staining (Fig. 1B). In FACS analysis, reduced granularity and smaller size of PMC-306 was confirmed. The population of primary WT PMCs shifted towards higher SSC intensities and showed more heterogeneity compared to PMC-306 (Fig. 1C). However, though differences appeared in morphological characteristics, PMC-306 stained positive for the typical MC markers FcεRI, KIT, ST2, and CD13 (Fig. 1D, suppl. Fig. 1B). Remarkably, KIT expression at the cell surface was 10-fold increased, when compared to primary WT PMCs, while expression of other surface markers (FcεRI, ST2 and CD13) was not significantly altered (Fig 1D, suppl. Fig. 1C) suggesting an important pro-survival role for KIT in the PMC-306 cell line.

### PMC-306 show increased but still cytokine-dependent proliferation

Established human MC leukemia cell lines like HMC-1.1 and HMC-1.2 are known for their high proliferative capacity and cytokine-independent growth due to mutations in the receptor-tyrosine kinase KIT (13, 35). Therefore, we sought to determine the cytokine dependence and proliferative capacity of PMC-306. We first compared the proliferation rate of PMC-306 to primary PMCs over 3 days in SCF- and IL-3-containing culture medium. PMC-306 had a doubling time of about 24 hours while primary PMCs proliferated marginally within an investigated time frame of 72 h (Fig. 2A). Due to the slow proliferation of primary PMCs, we analyzed the impact of IL-3 and SCF withdrawal on the proliferation only for PMC-306 cells. Both cytokines were reduced independently, while one factor was kept constant. Reduction of IL-3 to 1% and complete withdrawal of IL-3 significantly, but moderately reduced proliferation after 72 hours, while at earlier time points proliferation remained unaffected (Fig. 2B). Instead, reducing the amount of SCF had a substantial impact on proliferation of PMC-306. Reduction by 90% substantially attenuated proliferation already at 48 hours after withdrawal. Further reduction of SCF completely abrogated proliferation of PMC-306 (Fig. 2C). Additionally, we measured the viability of primary WT PMCs and PMC-306 after 72 hours of incubation under different deprivation conditions. Reducing the amount of IL-3 had no impact on viability of primary PMCs but significantly attenuated the viability of PMC-306 (Fig. 2D, suppl. Fig. 2). Reduction of the SCF concentration significantly reduced viability of primary PMCs. However, 10% remaining SCF reduced viability of PMC-306 without affecting viability of WT PMCs. Under harsher SCF deprivation conditions, viability of PMC-306 cells decreased stronger indicating increased SCF dependence of PMC-306 cells over WT PMCs (Fig. 2D, suppl. Fig. 2A/B). Interestingly, when metabolic activity under IL-3-deprivating conditions was measured, primary PMCs maintained higher metabolic rates compared to a faster decline in metabolic activity of PMC-306 cells (Fig. 2F). SCF withdrawal reduced metabolic activity similarly in both cell types (Fig. 2F). In line with increased KIT expression and stronger dependence on SCF for survival, PMC-306 cells were more sensitive to KIT inhibition by Imatinib. 1 µM of Imatinib was sufficient to significantly reduce metabolic activity in PMC-306 cells while primary PMCs appeared more resistant showing sustained metabolic activity at the same inhibitor concentration (Fig. 2G). Conclusively, the PMC-306 cell line is slightly dependent on IL-3, and even more – compared to primary WT PMCs – on SCF for proliferation and survival.

**Fig. 2:**
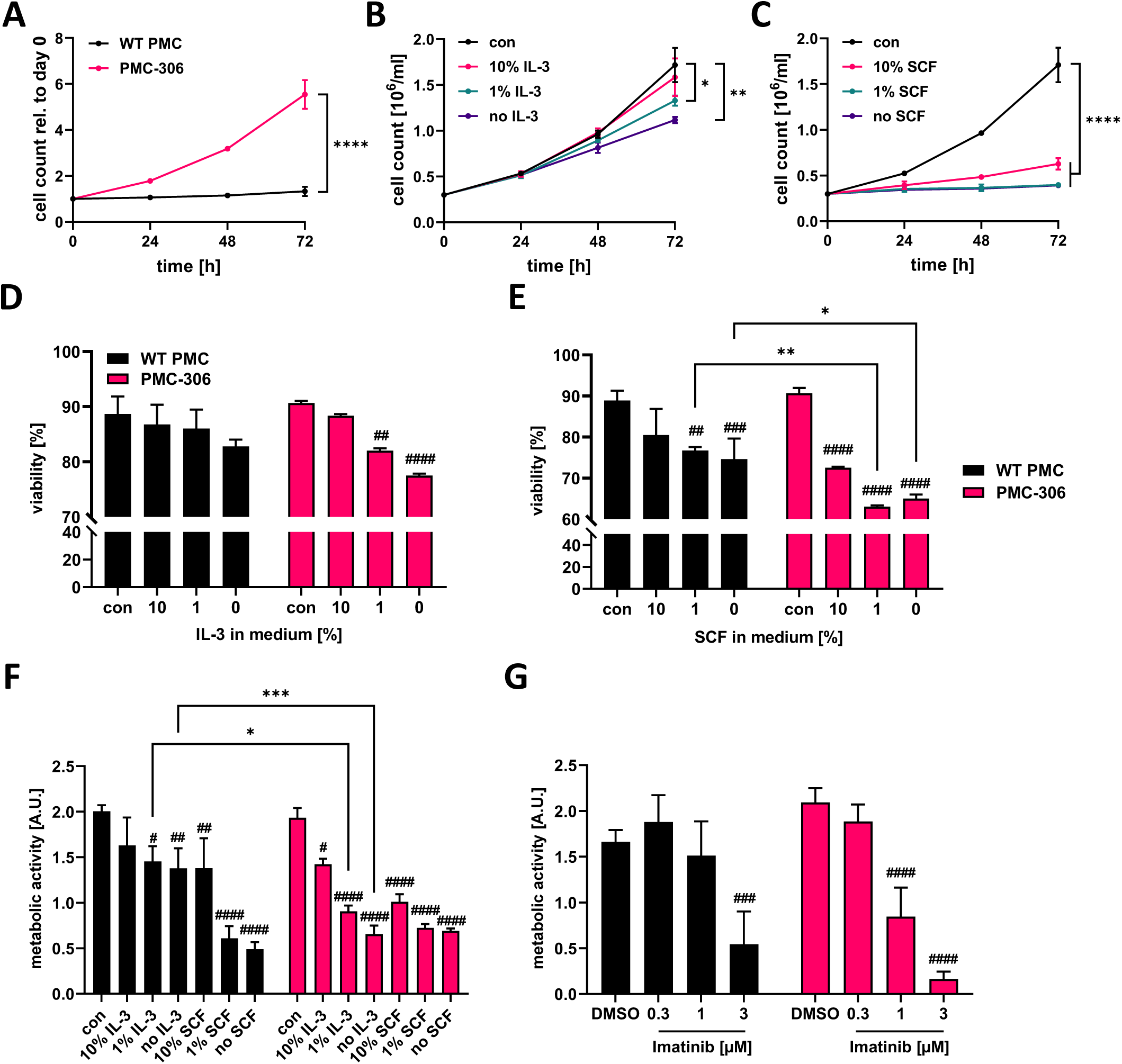
PMC-306 show faster proliferation compared to primary PMCs, which is dependent on IL-3 and SCF. **(A)** Cell number of primary WT PMCs and PMC-306 cells was measured every 24 hours for up to 72 hours using a CASY Cell counter (n=3). Proliferation of PMC-306 cells grown under reduced IL-3 (n=3) **(B)** or SCF (n=3) **(C)** concentrations was determined every 24 hours up to 72 hours using a CASY Cell counter. Viability of primary PMCs and PMC-306 cells was determined by FACS analysis using Annexin V and propidium iodide after 72 hours incubation in medium containing reduced IL-3 (n=3) **(D)** or SCF (n=3) **(E). (F)** The metabolic activity of primary PMCs and PMC-306 after 72 hours cultivation under IL-3 or SCF deprivation was determined by XTT assays (n=3). **(G)** The metabolic activity of Imatinib-treated primary PMCs and PMC-306 cells was determined after 72 hours by XTT assays (n=3). Data are shown as mean + SD. **(A)** Two-tailed, unpaired Student’s T test of data at 72 hours. **(B,C)** One-way ANOVA followed by Tukey multiple comparisons test of data at 72 hours. **(D)-(G)** Ordinary two-way ANOVA followed by Sídák multiple comparisons test. *p*>0.05 ns, # and * *p*<0.05, ## and ** *p*<0.01, ### and *** *p*<0.001, #### and **** *p*<0.0001. * indicates significant differences between groups, while # indicates significant differences relative to control conditions within one group.

### Ag-dependent MC effector functions of PMC-306 are increased

Leukemic MCs proliferate fast, while the reaction to stimulatory signals associated with degranulation and cytokine production is often attenuated due to their immature phenotype or loss of receptor expression (e.g. the FcεRI α- and β-subunits in case of HMC-1.1 and -1.2 cells) (36–38). Therefore, we were interested in determining the capacity of degranulation and pro-inflammatory cytokine production of PMC-306 cells in response to different stimuli. The FcεRI is known to respond to increasing concentrations of Ag with a bell-shaped dose response curve (39). The activity of released β-hexosaminidase from cytoplasmic granules can be used as readout for degranulation. Alternatively, to the latter bulk measurement, externalization of the granular transmembrane marker LAMP1 can be measured by FACS analysis to quantify the extent and kinetics of granule fusion with the plasma membrane on a single cell basis. PMC-306 and primary PMCs both showed similar bell-shaped dose-response behaviour with increasing Ag concentrations, reaching a maximum release of β-hexosaminidase at 20 ng/ml (Fig. 3A). Additionally, we determined the kinetics of degranulation measured as time-dependent LAMP1 externalization. Interestingly, in PMC-306 cells LAMP1 externalization was stronger than in primary PMCs suggesting enhanced granule membrane fusion on a single cell basis. Yet, the kinetics of LAMP1 externalization was similar reaching maximal LAMP1 MFIs after 5 minutes of stimulation (Fig. 3B). Another receptor triggering MC degranulation independent of IgE is the GPCR MRGPRB2, which is exclusively expressed in connective tissue-like MCs and can be stimulated by cationic peptides (40–42). Thus, to prove expression of MRGPRB2 in PMC-306, we stimulated primary PMCs and the PMC-306 line with either C48/80 or Mastoparan – compounds that are known to potently activate MRGPRB2 – and measured β-hexosaminidase release. Primary PMCs and PMC-306 both de-granulated to a similar extent in response to both stimuli in a dose-dependent manner confirming the connective tissue nature of PMC-306 (Figs. 3C & D). While degranulation of MCs happens within minutes, secretion of pro-inflammatory cytokines takes several hours of stimulation. We determined secretion of IL-6 and TNF from primary PMCs and PMC-306 in response to FcεRI stimulation. As described above, the response to Ag stimulation followed a bell-shaped dose-response curve in both cell types (Fig. 3E). Intriguingly, PMC-306 secreted significantly higher amounts of IL-6 and TNF compared to primary WT PMCs across all Ag concentrations tested. Besides, the threshold for cytokine release was lower for PMC-306 as a concentration of 0.2 ng/ml of Ag was sufficient to trigger detectable cytokine production. We further observed no pro-inflammatory cytokine secretion in response to TLR4 (LPS), TLR2/6 (FSL-1), KIT (SCF) and MRGPRB2 (C48/80) stimulation at the given stimuli concentrations for both cell types (Fig 3F). However, primary PMCs and PMC-306 reacted to IL-33 activating the ST2/IL1RAP (IL-1 receptor accessory protein) receptor complex. The response of PMC-306 cells to IL-33 was significantly diminished although the ST2 receptor expression levels at the cell surface were similar to primary PMCs (Fig. 3F, Fig. 1C). Briefly, we could show that the PMC-306 line expresses a functional FcεRI and MRGPRB2 receptor, maintained the ability to degranulate in response to Ag and MRGPRB2 activation, and exhibited a hyperactive phenotype regarding IL-6 and TNF production in response to Ag stimulation, while the IL-33-triggered cytokine production was reduced.

**Fig. 3:**
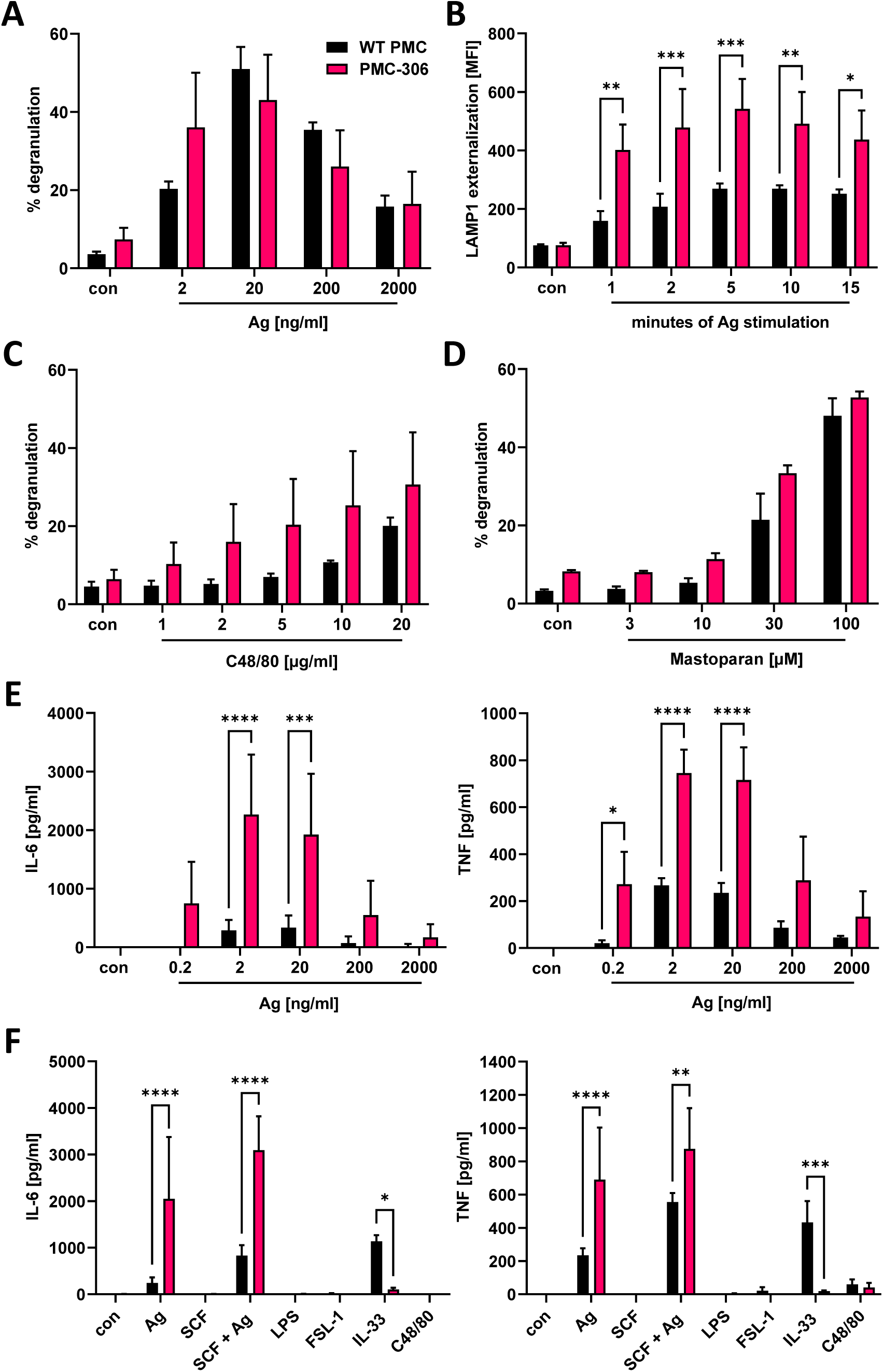
The qualitative degranulation and proinflammatory cytokine responses of PMC-306 are similar, albeit quantitatively stronger compared to primary WT PMCs. **(A)** Degranulation of primary WT PMCs and PMC-306 cells in response to increasing Ag concentrations was determined by β-hexosaminidase release assay (n=3). **(B)** The time-dependent externalization of LAMP-1 was determined by quantifying the LAMP-1 on the cell surface in FACS analysis as MFI upon stimulation with 20 ng/ml Ag (n=3). The degranulation responses of primary PMCs and PMC-306 cells to increasing concentrations of the MRGPRB2 agonists C48/80 **(C)** and Mastoparan **(D)** were determined by β-hexosaminidase release assays (n=3). The secretion of IL-6 **(E, left)** and TNF **(E, right)** in response to indicated Ag concentrations was determined by ELISA (n=3). The pro-inflammatory cytokine production of primary WT PMCs and PMC-306 to Ag (20 ng/ml) in comparison to other stimuli (SCF: 100 ng/ml, SCF+Ag: 100 ng/ml and 20 ng/ml, LPS: 1µg/ml, FSL-1: 1 µg/ml, IL-33: 10 ng/ml, C48/80: 10 µg/ml) was determined by IL-6 **(F, left)** and TNF **(F, right)** ELISAs (n=3). Data are shown as mean +SD. Two-way ANOVA followed by Sídák multiple comparisons test. *p*>0.05 ns, * *p*<0.05, ** *p*<0.01, *** *p*<0.001, **** *p*<0.0001.

### Signaling pathway activation downstream of typical MC receptors is functional in PMC-306 cells

In comparison to BMMCs, PMCs have a more aggressive protease composition stored within the cytoplasmic granules. This complicates signaling studies and makes protein interaction studies using immunoprecipitation nearly impossible as the lysis of cells in classical lysis buffers containing a mixture of protease inhibitors is not sufficient to prevent protease-dependent protein degradation (12,34,43). We therefore aimed to test whether the PMC-306 is suitable for signaling studies upon lysis in normal lysis buffer containing typical protease inhibitors. Further, we wanted to confirm that differential activation of the PMC-306 line leads to comparable activation of signaling pathways as in primary PMCs to verify its suitability for MC signaling studies. Stimulation of PMC-306 with Ag rapidly increased phosphorylation of PLCγ1 at tyrosine 783, which is critical for the initiation of the Ca^2+^ response and PKC activation. Activation of the mitogen-activated protein kinases (MAPK) ERK1/2 and p38, as well as phosphorylation of protein kinase B (PKB) occurred within one minute of stimulation, while strongest phosphorylation/activation and degradation of NFκB inhibitor α (IκBα) – a readout for NFκB pathway activation – started at 5 minutes after Ag stimulation (Fig. 4A). This indicates that pathways necessary for cytokine production, like the MAPK and NFκB pathways, are functional and strongly responsive to Ag stimulation, which is in line with the described cytokine measurements (Figs. 3E & F). Although we could not observe cytokine secretion in response to SCF stimulation, activation of KIT (auto-phosphorylation at Y719) could be detected in response to SCF. However, phosphorylation of KIT, PKB, and MAPK were transient and rapidly decreased after one minute of stimulation (Fig. 4B) correlating with the lack of detectable cytokine production, which would require a sustained signal strong enough to induce transcriptional changes in gene expression. Likewise, we could see transient activation of the NFκB and ERK pathways in response to IL-33 though the cytokine production was strongly attenuated compared to WT PMCs (Fig. 3F). Interestingly, Y719 of KIT was phosphorylated upon IL-33 stimulation, which corroborates previous findings that revealed a role for KIT in contributing to ST2 signaling in MCs (Fig. 4C) (44). Additionally, while most cell lines have a strong basal activation of pro-survival pathways due to oncogenic mutations, we did not observe obvious activation of KIT, PKB or ERK in PMC-306 cells. Surprisingly, we could not detect considerable cytokine production in response to C48/80 at a concentration of 10 µg/ml, which in previous experiments elicited substantial β-hexosaminidase release (Fig. 3C) and was also used in preceding studies to stimulate MRGPRB2 (40). Yet, we noticed a strong induction of cytokine release in response to 3-fold lower C48/80 concentrations (suppl. Fig 3A). Accordingly, we used 3 µg/ml C48/80 to analyze activation of MRGPRB2 signaling in the PMC-306 cell line. As shown in Fig. 4D, a transient activation of the PKB, ERK and NFκB signaling pathways could be detected after five minutes of stimulation. Notably, the concentrations stimulating cytokine production did not or did only hardly induce degranulation. Besides, stimulation with higher concentrations of C48/80 and Mastoparan previously reported to induce degranulation led to a significant reduction in protein concentration in the cell pellet after lysis, which did not occur with Ag (suppl. Fig. 3B), indicating loss of cellular content upon C48/80 stimulation. Moreover, we were not able to measure an externalization of LAMP1 in response to C48/80 at 10 µg/ml neither in PMC-306 cells (data not shown) nor in primary PMCs (suppl. Fig. 3C) pointing to a substantially different mechanism of degranulation compared to Ag, which potentially involves cellular disintegration.

**Fig. 4:**
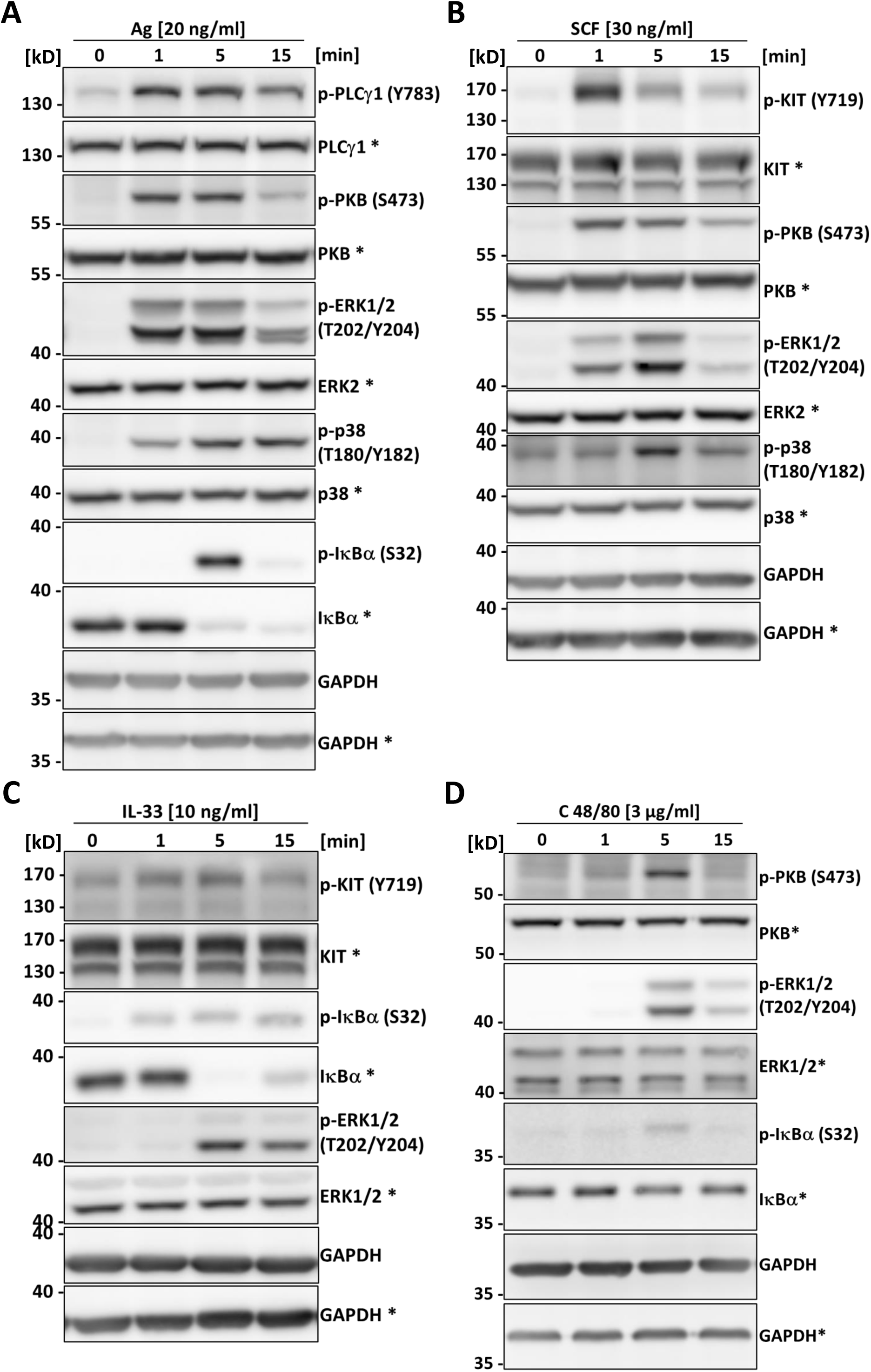
The PMC-306 cell line can be used to analyze signaling in response to Ag, SCF, IL-33 and C48/80. The activation of PLCγ1, PKB, the MAPKs ERK and p38, KIT and the NFκB pathway in response to Ag (20 ng/ml) **(A)**, SCF (30 ng/ml) **(B)**, IL-33 (10 ng/ml) **(C)** or C48/80 (3 µg/ml) **(D)** after 1, 5 and 15 minutes of stimulation was studied by Western blot analysis. Activation of the pathways was analyzed by phospho-specific antibodies (p-PLCγ1 (Y783), p-KIT (Y719), p-PKB (S473), p-ERK1/2 (T202/Y204), p-p38 (T180/Y182), p-IκBα (S32) for key signaling proteins. For loading controls, antibodies recognizing the respective total proteins were used. Asterisks indicate detection on the same membrane. GAPDH served as control for equal loading of both gels.

### Reduced protease expression of PMC-306 suggests an immature phenotype

Our data so far demonstrated that the PMC-306 line still has MC-like properties in terms of morphology and cellular reactivity. Nonetheless, we also noted differences in proliferation, a higher cytokine secretion and the fact, that in contrast to primary WT PMCs, PMC-306 has become immortal. Hence, we were interested in transcriptomic differences between PMC-306 and primary WT PMCs that we aimed to uncover by next generation sequencing. For validation of NGS data, we first concentrated on variations in MC granular proteins as the electron micrographs revealed differences in number of granules between WT PMCs and PMC-306. We determined the relative β-hexosaminidase content of equal cell numbers of primary PMCs and PMC-306 and found significantly less β-hexosaminidase activity in cell lysates of PMC-306 (Fig. 5A). On mRNA level, we found that expression of the proteases *Cpa3*, *Cma1* and *Gzmb* was reduced compared to WT PMCs but still detectable in PMC-306 (Figs. 5B-D). However, we could not detect expression of *Mcpt2*, *Mcpt4* and *Tpsab1* in PMC-306 cells (Figs. 5E-G). On protein level, we verified expression of granzyme B and tryptase in primary WT PMCs and could confirm a total lack of tryptase expression in PMC-306 cells. Strikingly, despite residual mRNA expression of *Gzmb* in PMC-306, there was hardly any granzyme B protein present (Fig. 5H). This pattern of protease expression in PMC-306 is reminiscent of immature MC precursors, that lack expression of proteases up-regulated in late stages of MC differentiation like *Mcpt2* and *Tpsab1* (45, 46). In addition, we wanted to verify expression of the CTMC-specific receptor MRGPRB2, since we could show similar responsivity between PMC-306 and WT PMCs (Figs. 3C & D). Though, PMC-306 cells exhibited significantly reduced mRNA expression of *Mrgprb2* (Fig. 5I). However, our data suggests that this does not affect the respective degranulation response (compare Fig. 3C).

**Fig. 5:**
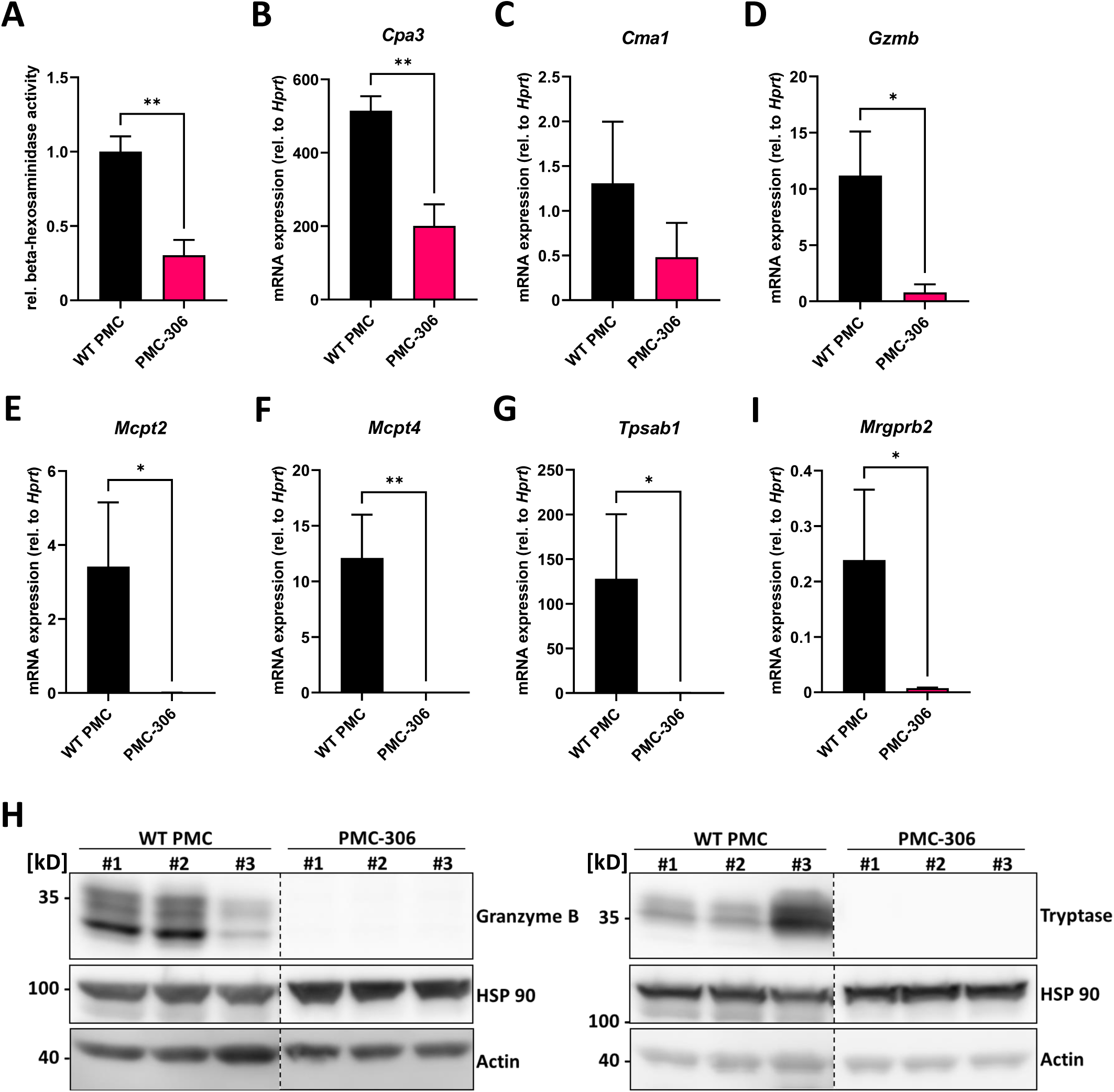
The repertoire of protease expression is reduced in PMC-306 cells revealing an immature MC phenotype. **(A)** The β-hexosaminidase content in primary PMCs and PMC-306 cells was determined by measuring the relative β-hexosaminidase activity in equal numbers of primary PMCs and PMC-306 cells (n=3). **(B)-(G)** The protease expression of *Cpa3*, *Cma1*, *Gzmb*, *Mcpt2*, *Mcpt4* and *Tpsab1* (*Mcpt7*) and the expression of *Mrgprb2* **(H)** were quantified by RT-qPCR in samples from fully differentiated primary PMCs and PMC-306 cells. Data are expressed as mRNA expression relative to *Hprt* determined according to the deltaC_t_ method (n=3). **(H)** Representative Western blots of three lysates from three independent primary WT PMCs and independently taken samples from PMC-306 cells comparing the expression of granzyme B and tryptase on protein level. HSP 90 and actin served as loading controls. Data are shown as mean +SD. Unpaired, two-tailed Student’s *t*-test with Welsh’s correction. * *p*<0.05, ** *p*<0.01.

### PMC-306 cells are characterized by a loss of *Cdkn2a* and *Arf* tumour suppressor gene expression

Next, we addressed the question, which genomic or transcriptional alterations are responsible for the transformation and increased proliferation of the PMC-306 line. We first analyzed the karyotype of PMC-306 cells, which revealed a typical male murine diploid (19XY) telocentric set of chromosomes (Fig. 6A). However, a prominent structural aberration of chromosome 4 characterized by a heterozygous interstitial deletion (Del(4)(C4-C7) was detected (Fig. 6A, marked by a red arrow). This structural aberration was consistent in all analyzed cells (suppl. Fig. 4A) as depicted in Fig. 6B showing three further examples of partial karyotypes of the heterozygously deleted chromosome 4 region (Del(4)(C4-C7). The second most prominent chromosomal aberration represented a Y-autosome translocation (T(Y;17)) (Fig. 6C) appearing in roughly one third of analyzed karyotypes (suppl. Fig. 4A), which has not been described so far so that its significance for the transformation process is presently unclear.

**Fig. 6:**
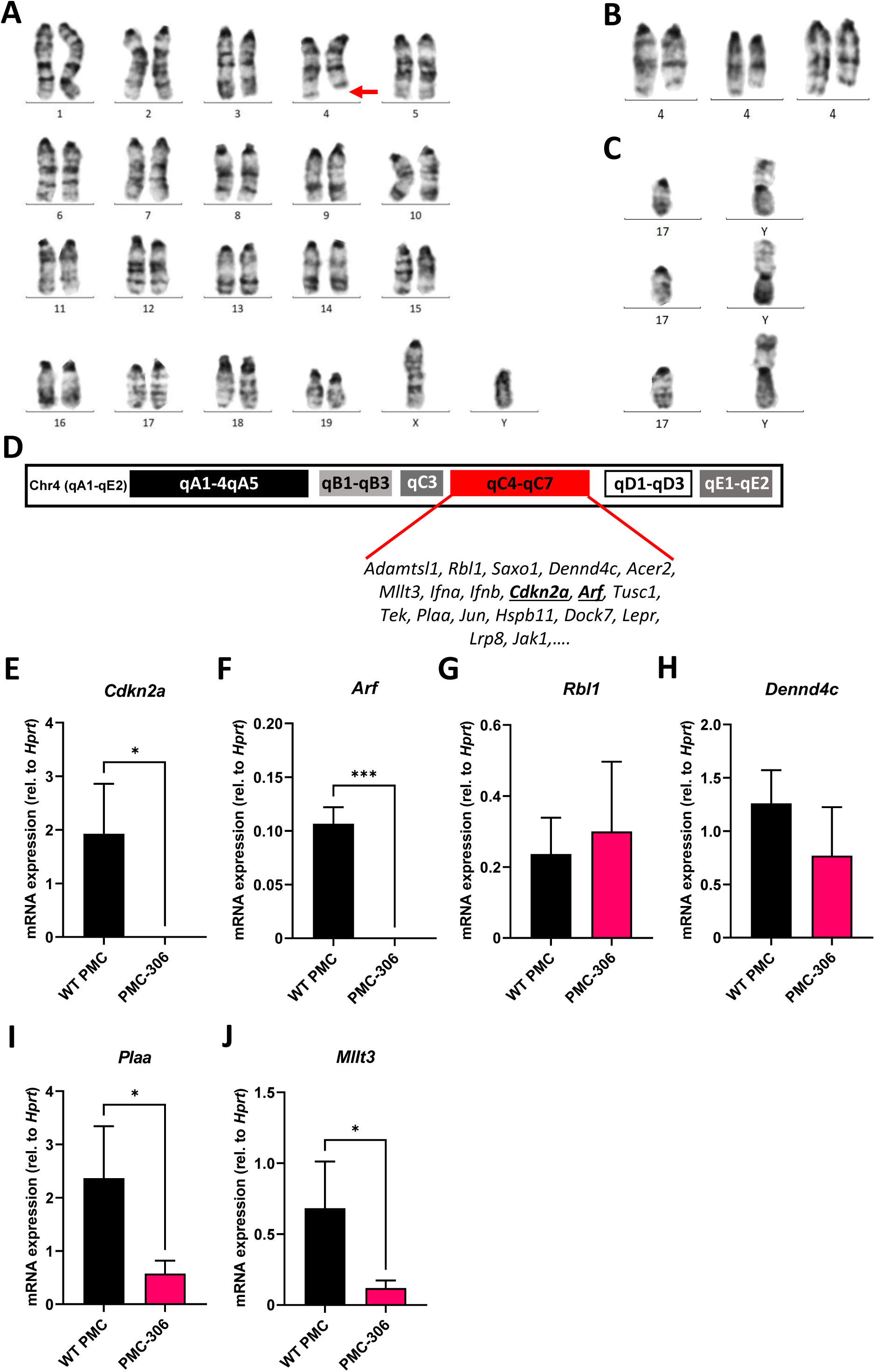
Cytogenetic and gene expression analysis identified loss of *Cdkn2a*/*Arf* expression as potential cause for immortalization of PMC-306. Mitotic chromosomes of PMC-306 were cytogenetically analyzed. **(A)** Representative image of a conventional G-banded karyotype with structural aberration. Red arrow indicates interstitial deletion of chromosome 4 (Del(4)(C4-C7)). **(B)** Further examples of partial karyotypes showing structurally aberrant chromosomes 4 with the same interstitial deletion. **(C)** Three partial karyotypes representing the Y-autosome translocation (T(Y;17)) – confirmed by fluorescence in situ hybridization with a Y-specific whole mouse chromosome painting-probe (confirmatory FISH analysis not shown)). **(D)** Architecture of murine chromosome 4 with highlighted region qC4-qC7 and exemplary genes encoded within this region. **(E)-(J)** RT-qPCR analysis of *Cdkn2a*, *Arf*, *Rbl1*, *Dennd4c*, *Plaa* and *Mllt3* mRNA expression in primary WT PMCs and PMC-306 cells. Data are expressed as mean +SD. Unpaired, two-tailed Student’s t test with Welsh’s correction. * *p*<0.05, *** *p*<0.001.

Interestingly, within the heterozygously deleted region of chromosome 4 the INK4/ARF locus is encoded (Fig. 6D) (47). p16/INK4A and p19/ARF are responsible for regulation of the RB and p53 pathways, respectively, thus controlling two essential cell cycle regulators (25, 48). As PMC-306 cells showed a faster proliferation than WT PMCs, we suspected that expression of genes within the INK4/ARF locus might be reduced in PMC-306 cells. We could confirm that mRNA expression of *Cdkn2a* encoding INK4A and *Arf* was not only reduced – as one would expect as a result of a heterozygous deletion – but completely absent in PMC-306 cells (Figs. 6E & F). Remarkably, other genes located in the same chromosomal region exhibited a different behaviour. Expression of *Rbl1* and *Dennd4c* was unaffected suggesting that one allele is sufficient to compensate for the deletion of the second (Figs. 6G & H), while expression of *Plaa* and *Mllt3* was reduced compared to primary PMCs as one could expect if one allele is lost (Figs. 6I & J). Curiously, an independently mutated PMC line, PMC-303, not showing the interstitial deletion on chromosome 4 (suppl. Fig. 4B) also completely lost *Cdkn2a* and *Arf* expression (suppl. Figs. 4C & D), while other genes within the same chromosomal region were not (*Rbl1*, *Mllt3*, *Plaa*) or only slightly (*Dennd4c*) affected (suppl. Figs. 4E-H). In addition, PMC-303 cells showed a trisomy 8 due to a homologous Robertsonian translocation of chromosome 8 (Rb(8.8)) in 24 of 35 analyzed metaphasic spreads (suppl. Fig. 4B), which we did not observe in PMC-306 cells.

### The PMC-306 cell line has a de-regulated cell cycle

It is well known that uncontrolled proliferation and loss of cell cycle regulation is one of the hallmarks of cancer (25,48–51). As we have shown above, PMC-306 cells have a strongly accelerated proliferation, which likely is a consequence of the observed loss of *Cdkn2a* and *Arf* expression. Hence, we supposed that in PMC-306 the cell cycle is de-regulated to enable fast proliferation. To prove this, we performed RNAseq with both PMC lines PMC-303 and -306 as well as primary WT PMCs as control. RNA expression was analyzed from cells directly taken from conventional culture medium without additional treatment. We searched the mRNA expression data for global transcriptomic changes between primary PMCs and the PMC-306 line and focussed on genes that were at least 5-fold regulated. GO enrichment analysis unveiled that the top 15 GO terms were all related to cell cycle, mitosis and cell division (suppl. Fig. 5A). This result led us to investigate the cell cycle by comparing the obtained long-term cultures of spontaneously transformed PMC-306 line with primary WT PMCs using flow cytometry. The gating strategy and unstained controls for background determination for each cell type are shown in supplementary Figs. 5B-D. We first examined cell cycle activity in non-transformed primary WT PMCs and the PMC-306 cell line using the established general cell cycle marker MKI67. As expected, PMC-306 cells revealed substantially (*i.e.* approximately twofold) higher cell cycle activity when compared to primary WT PMCs in terms of MKI67 positive cells (∼60-63% vs. 31%) and it’s expression strength as determined by mean fluorescence intensity (Figs. 7A & B). In good agreement, we further detected significantly enhanced mitotic activities in the PMC-306 cells in relation to primary WT PMCs after analyzing phospho-Histone H3 (pH3). PMC-306 cells displayed a higher amount of pH3 positive cells (2%) compared to primary PMCs (1%, Fig. 7C). Additionally, we determined the DNA content by DAPI-staining. As expected, MKI67 was found to be expressed in the cell cycle phases Gap1 (G_1_, 2n), DNA-synthesis (S-phase, 2-4n) to Gap2 (G_2_, 4n) and mitosis (M-phase, 4n) in primary WT PMCs and PMC-306 cells (suppl. Fig. 5E, left panel). Furthermore, mitotic active pH3-positive cells were specifically found at a DNA content of 4n in primary PMCs and PMC-306 cells (suppl. Fig. 5E, right panel). Importantly, PMC-306, but not primary PMCs, additionally exhibited an MKI67 and pH3 positive cell fraction with a DNA content above 4n, indicating chromosomal abnormalities such as aneuploidy after the transformation step (suppl. Fig. 5E). Finally, we investigated the DNA integrity by staining of phospho-histone 2Ax (pH2Ax), a well-described marker for DNA-double strand breaks. We found low levels of pH2Ax in WT PMCs (∼1%) and slightly increased levels in PMC-306 (∼2%, Fig. 7D). Together, the cell cycle of PMC-306 cells is characterized by a higher mitotic activity, increased DNA damage and aneuploidy.

**Fig. 7:**
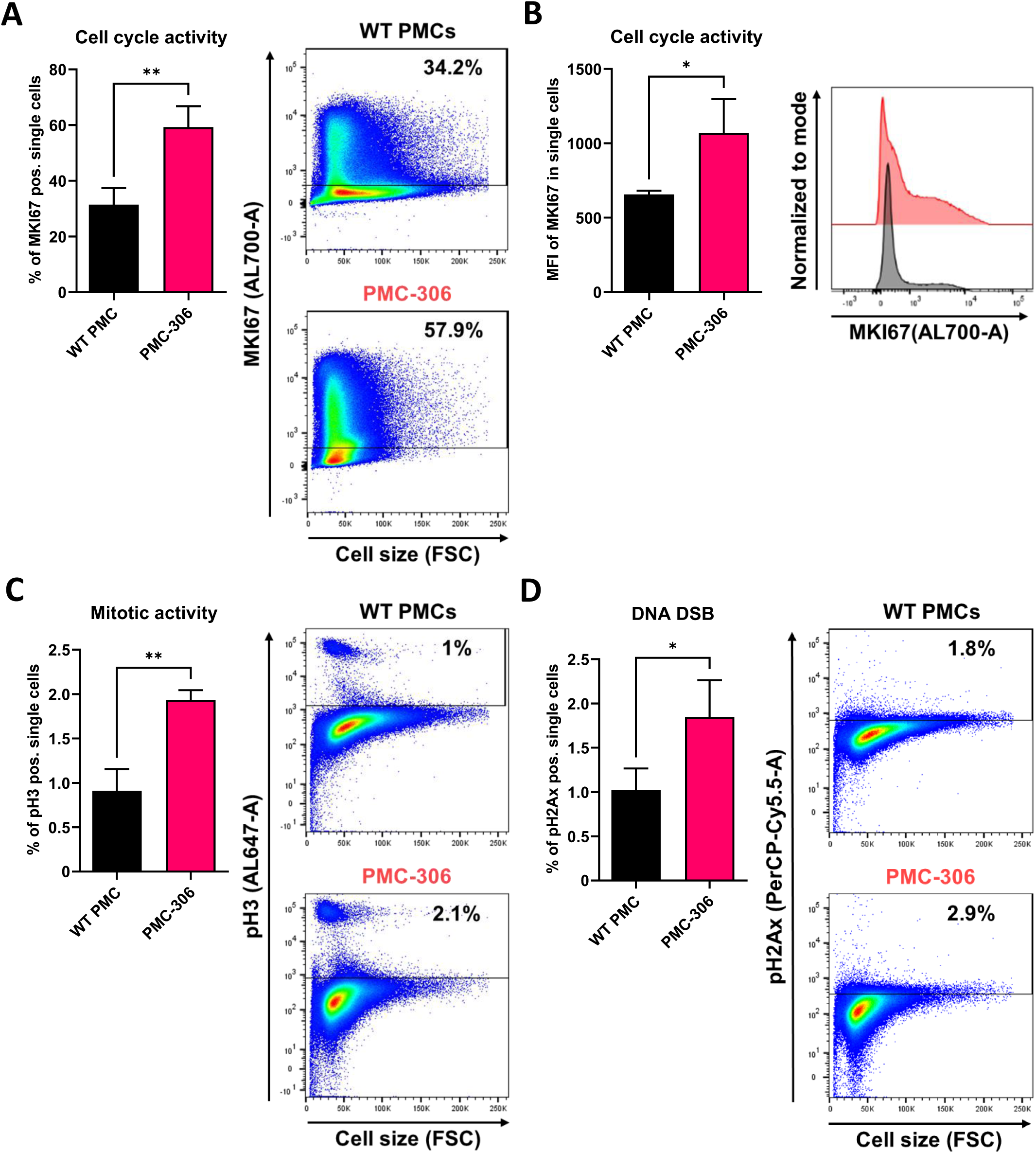
PMC-306 cells are characterized by enhanced cell cycle activity and sporadic DNA damage. The cell cycle of primary WT PMCs was compared to PMC-306 cells. **(A)** Cells were stained with an antibody directed against MKI67 (AL700-A). Left: Quantification of MKI67+ cells (% of single cells, n=3). Right: Representative FACS-plots of MKI67+ cells (% of single cells). **(B)** Quantification of the mean fluorescence intensity (MFI, n=3) of MKI67 in single cells (left) and a representative histogram showing MKI67 intensities in primary PMCs and PMC-306 cells. **(C)** Cells were stained with an antibody directed against phosphorylated histone H3 (pH3, AL647-A). Left: Quantification of pH3 cells (% of single cells, n=3). Right: Representative FACS-plots of pH3+ cells (% of single cells). **(D)** Cells were stained with an antibody directed against phosphorylated Histone H2Ax (pH2Ax, PerCP-Cy5.5-A). Left: Quantification of pH2ax cells (% of single cells, n=3). Right: Representative FACS-plots of pH2Ax+ cells (% of single cells). Data are expressed as mean +SD. Unpaired, two-tailed Student’s t test with Welsh’s correction. * *p*<0.05, ** *p*<0.01.

### Constitutive KIT activation leads to loss of *Cdkn2a* and *Arf* expression in BMMCs

Finally, we aimed at identifying the mechanism for loss of *Cdkn2a/Arf* expression in primary PMCs and suspected KIT activation as a crucial pro-survival and potentially transformation-promoting mechanism due to the occurrence of frequent KIT mutations in leukemic MC lines (35, 52). Therefore, we analyzed the behaviour of BMMCs, which do not necessarily need SCF for proper differentiation into MCs, and cultivated them from the beginning of the differentiation phase in the absence or presence (mimicking PMC culture conditions) of SCF. We noticed remarkable phenotypic changes of BMMC cultures induced by SCF supplementation. After the differentiation phase of 4 weeks, BMMCs cultivated in the presence of SCF were larger (increased FSC) and more granular (increased SSC) compared to BMMCs from the same mouse only supplemented with IL-3 (Fig. 8A left two panels, Fig. 8B left and middle, Fig. 8C left and middle). Intriguingly, these morphological differences disappeared after 10 weeks, when size and granularity of SCF-supplemented BMMCs decreased to levels of BMMCs cultivated without SCF (Fig. 8A right panels, Fig. 8B and C). As a control for SCF treatment we stained for KIT at the cell surface. KIT is known to be internalized upon SCF binding (53). Consequently, less KIT remains on the cellular surface under chronic KIT-stimulating conditions. We could confirm this observation by FACS analysis, as BMMCs supplemented with SCF expressed significantly less KIT at the plasma membrane (Fig. 8B right panel, Fig. 8C right panel). It is further known that KIT can increase proliferation by inducing transcription of cyclins *via* the PI3K-AKT pathway (54). Indeed, we observed increased proliferation of SCF-treated BMMCs and strong upregulation of the G_1_/S cyclin *Ccnd1* until 8 weeks. After 10 weeks, however, *Ccnd1* expression decreased constantly to levels similar to BMMCs grown in the absence of SCF (Fig. 8D, left). Notably, we found a strikingly similar expression pattern for the negative cell cycle regulator *Cdkn2a*, whose expression was increased by SCF supplementation over the first 8 weeks and then dropped to levels below the expression of control BMMCs (Fig. 8D, middle). Accordingly, expression of *Arf*, though not significantly increased in the beginning, drastically decreased after 10 weeks in the presence of SCF, as well (Fig. 8D, right). Loss of *Cdkn2a* and *Arf* expression thus timely and reproducibly correlates with cell shrinkage and perfectly mimics the phenotype of PMC-306 cells. Hence, this observation suggests that constitutive KIT activation might be the driver for cell cycle de-regulation and subsequent MC transformation.

**Fig. 8:**
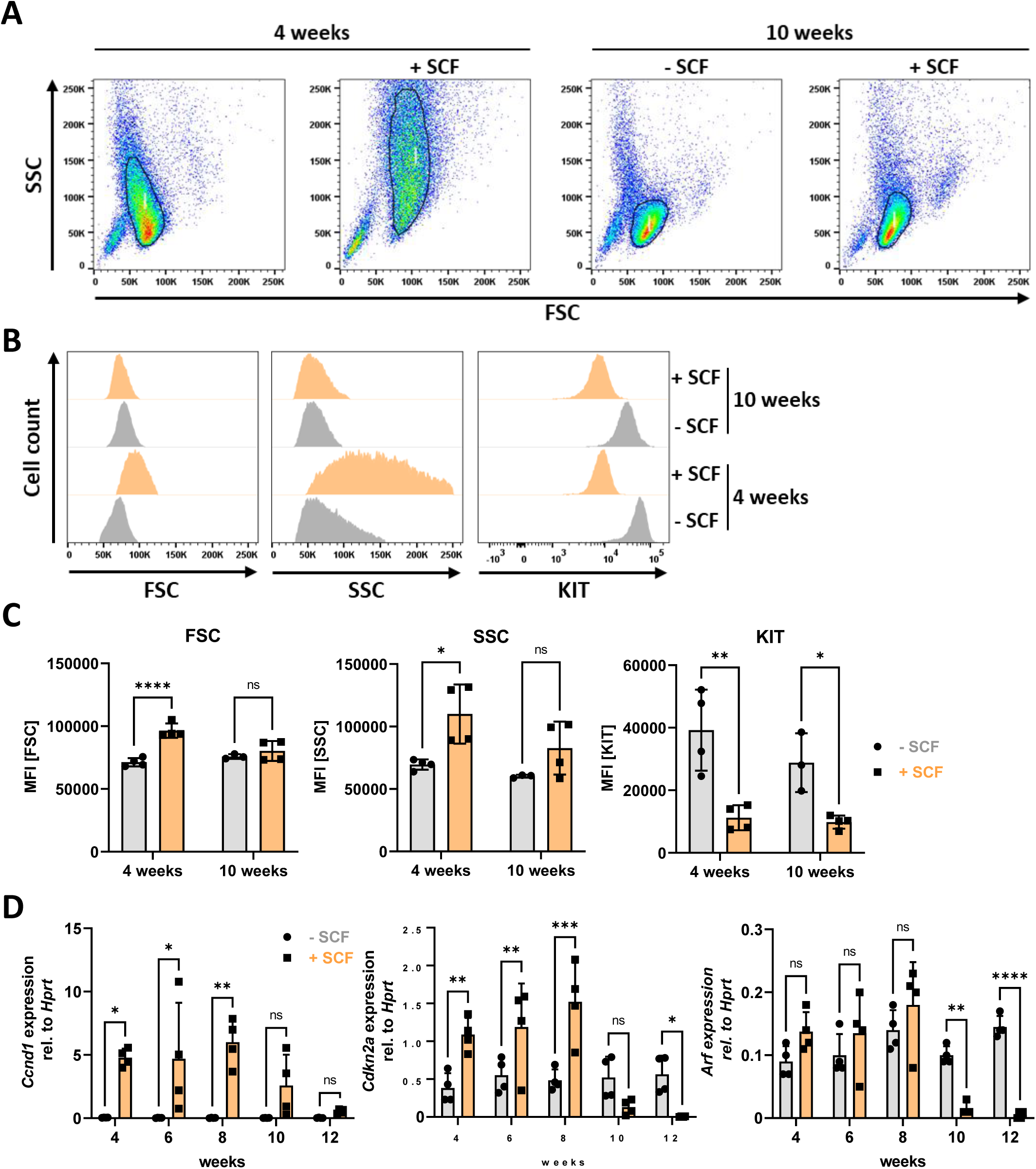
Constitutive KIT activation induces morphological changes and loss of *Cdkn2a/Arf* expression in BMMCs. **(A)** Representative FACS-plots showing BMMC populations cultivated with or without SCF for 4 (left) or 10 (right) weeks. **(B)** Representative histograms showing frequency distributions of forward scatter (left), side scatter (middle) and KIT expression (right) of BMMCs cultivated for 4 or 10 weeks with or without SCF. **(C)** Quantified data (MFI) of parameters analyzed in **(B)**. **(D)** RT-qPCR analysis of *Ccnd1*, *Cdkn2a* and *Arf* mRNA expression in BMMCs supplemented or not with SCF over a period of 12 weeks. Symbols indicate biological replicates. **(C)** and **(D)** Ordinary two-way ANOVA followed by Sídák multiple comparisons test. *p*>0.05 ns, * *p*<0.05, ** *p*<0.01, *** *p*<0.001, **** *p*<0.0001.

## Discussion

Advancing research of the characteristics and activation mechanisms of MCs either requires the isolation of MCs from donor tissue, *ex vivo* differentiation of CD34-positive precursor cells or the use of adequate cell lines. Problematically, suitable cell lines with MC-like properties are restricted to few examples with several limitations including loss of FcεRI expression in HMC-1.1, -1.2 and LUVA cells or long doubling times in case of LAD2 cells (14,15,36). Thus, MC research is encumbered by MC lines sharing as much features as possible with tissue-resident MCs *in vivo*. Here we present a novel murine MC line, which overcomes these limitations. We showed that the PMC-306 cell line stably expresses a functional FcεRI, has a short doubling time, maintained MC characteristics over multiple passages and is easy to freeze and thaw. These features suggest that PMC-306 cells can be used as a new tool to address questions related to MC research.

We extensively characterized PMC-306 cells and compared their appearance, activation and growth behaviour to non-transformed primary WT PMCs. PMC-306 cells share all characteristics of MCs as they stained positive for the MC surface markers FcεRI, KIT (CD117), CD13 and T1/ST2. We could further demonstrate a strictly IL-3- and SCF-dependent growth and Imatinib sensitivity of PMC-306 excluding KIT activating or Imatinib-resistant mutations, which are often detected in MC leukemia and complicate studies on general MC properties. HMC-1.1 and -1.2 cells bearing KIT mutations have a high basal activity of pro-survival signaling pathways including PI3K/PKB, STAT5 and RAS/RAF/MEK/ERK (52). This complicates studies on signaling pathways activated by other receptors due to enhanced background signals. Importantly, PMC-306 cells, despite cultivated in SCF-containing medium, did not show these high background phospho-signals making them an interesting tool to study MC signaling.

The PMC-306 cell line is derived from primary WT PMCs, which have connective tissue-like MC characteristics. We confirmed this by verifying expression and functionality of the connective tissue MC-specific receptor MRGPRB2. This MC-specific receptor has recently gained attention as its discovery has led to many new insights into the role of MCs in adverse drug reactions, communication with nerve cells and the development of a new tool for skin MC ablation (40,41,55). However, studies on the MC activating mechanisms in terms of signal transduction of MRGPRB2 are scarce. We showed that PMC-306 cells are an appropriate model to study – apart from FcεRI-dependent MC activation – also MRGPRB2-dependent effects. PMC-306 cells substantially reacted to MRGPRB2 activating ligands in terms of degranulation and cytokine production. Strikingly, the results showed that the degranulation seems to involve a completely different mechanism from that of Ag-triggered degranulation as we could not detect externalization of the granular marker LAMP-1 and cells significantly lost a bigger part of their protein content within the first minutes of stimulation. A study from Gaudenzio et al. (56) already proposed that MC activation *via* the MRGPRB2 receptor is faster, does not involve compound exocytosis of granules and that the cytokine secretion profile is substantially different in comparison to Ag-dependent degranulation. Nevertheless, they did not deeply look into a mechanistic level how this is regulated by the cell. We further determined the pro-inflammatory cytokine production potential of PMC-306 cells in response to different stimuli. Interestingly, both primary WT and PMC-306 cells potently responded to Ag and IL-33, while TLR agonists like LPS and FSL-1 did not elicit a response. This is in agreement with a proteomic study revealing that primary human connective tissue type MCs lack crucial innate immune receptors including TLR-2, -3, -4, -7, -8 and -11, which is conserved between mouse and human (41). Looking at the overall picture of MC effector functions, there was great similarity between primary WT PMCs and the PMC-306 line.

One hallmark of MCs is their numerous densely packed secretory granules within the cytoplasm storing preformed pro-inflammatory mediators. PMC-306 cells appeared less granulated, showed reduced total β-hexosaminidase content and lacked expression of *Tpsab1*, *Mcpt2*, *Mcpt4* and *Cma1*. However, proteases that appear early in MC differentiation like *Cpa3* and *Gzmb* (46) were still present on mRNA level, which might indicate that the culture of PMC-306 consists of an incompletely differentiated transformed connective tissue MC type. However, GZMB protein expression was entirely absent. This could be a consequence of the drastically increased cell cycle progression, which does not permit the cell to build up complex granular structures resulting in post-transcriptional attenuation of mRNA translation not essential for proliferation. Mechanistically, the lavage of the murine peritoneal cavity may lead to isolation of MCs of different differentiation stages including fully mature and immature MC types. Of note, malignant transformation of myeloid cells often occurs at early differentiation stages, where the proliferative potential is high (46,57,58). These circumstances might have contributed to the transformation of the PMC-306 cell line and would well fit to the observed phenotype.

Spontaneous transformation of cells *in vitro* can either occur due to activating mutations in pro-survival pathways and/or genetic loss of tumour suppressor proteins (49). Transformation of cells is often associated with loss of cell type-specific characteristics. To exclude potential effects of the immortalization on MC properties we aimed to address the reason for the transformation of PMC-306. We could exclude an activating mutation in the prominent MC growth factor receptor KIT. However, we found complete loss of expression of the two central cell cycle regulators *Cdkn2a* and *Arf* both located on the heterozygously deleted region of chromosome 4. While p16/INK4A inhibits CDK4 and CDK6, thereby preventing entry into S phase by activation of RB (59), p19/ARF inhibits the ubiquitin ligase MDM2, which leads to p53 stabilization and, amongst others, initiation of apoptosis (60, 61). Thus, absence of both of these proteins fuels the cell cycle, circumvents cell cycle control and accelerates cell proliferation as quantified for PMC-306 cells by MKI67 and pH3 staining. The rapid proliferation rate is likely accompanied by a reduced time to build up more complex intracellular structures like MC granules, which is in line with the reduced granular content of PMC-306. Importantly, we detected loss of *Cdkn2a* and *Arf* expression also in the independently transformed PMC-303 line, which we did not further characterize in this report. However, PMC-303 cells did not show any abnormalities on chromosome 4 suggesting a different mechanism of gene silencing potentially involving epigenetic mechanisms. Interestingly, the PMC-303 cell line had a trisomy 8, which frequently occurs in murine embryonic stem cells and is associated with increased proliferation, genome instability but does not seem to affect cellular differentiation (62, 63). Crucially, as we observed inactivation of the *Cdkn2a/Arf* locus in two different MC lines, this tumour suppressor locus seems to be a central player to prevent MC transformation and might also be interesting to investigate in the context of clonal MC diseases like mastocytosis or MC leukemia. So far, a *CDKN2A* mutation was only described in a case report of a patient suffering from MC leukemia with persistent myelodysplastic syndrome (64) without further characterization. As *Cdkn2a/Arf* inactivation appeared to be a recurrent phenomenon in MCs, we suspected that the culture conditions, notably the presence of SCF, might contribute to the transformation process. In this regard, it is known that activation of the epidermal growth factor receptor (EGFR) correlates with *Cdkn2a* loss in glioblastoma formation (65). Likewise, tyrosine kinase inhibitor resistance of Philadelphia chromosome-positive leukemia correlates with deletion of the *CDKN2* gene (66), and cooperativity between PTEN loss and loss of *Cdkn2a* and *Arf* in histiocytic sarcoma has been reported (67). This wealth of data suggests that activation of RTKs and downstream PI3K-PKB signaling, which induces cyclin expression and accelerates proliferation (54), is necessary for *Cdkn2a/Arf* inactivation. Indeed, we could show that chronic KIT activation by SCF downregulated *Cdkn2a* and *Arf* expression in BMMCs providing a mechanism for MC transformation *in vitro*. While SCF-supplemented BMMCs tried to counterbalance the enormous upregulation of *Ccnd1* by KIT activation through upregulation of *Cdkn2a* up to the first 8 weeks, prolonged SCF-treatment inactivated the negative regulatory cell cycle regulators. After the loss of *Cdkn2a/Arf* expression, high cyclin expression obviously became unnecessary because normal cyclin levels were sufficient to fuel the cell cycle in the absence of negative regulatory proteins. This could explain the decrease in *Ccnd1* expression at the same time expression of *Cdkn2a/Arf* is lost in SCF-treated BMMCs. In essence, KIT is an abundant RTK present on all types of differentiated MCs and a potent activator of the PI3K-PKB pathway, thus possessing all requirements to induce the above described cell cycle regulatory changes enabling unrestricted proliferation.

In summary, we herein characterized a new murine FcεRI- and MRGPRB2-positive connective tissue type-like MC line displaying similar characteristics to primary PMCs including several advantages. Technically, increased proliferation and reduced protease expression of PMC-306 overcomes the problems of cell number limitations and protein degradation in cell lysates using conventional cell lysis conditions (43) related to experiments with primary PMCs. Ultimately, alternatives to isolation of cells from and experiments with laboratory animals are needed to reduce the number of animals sacrificed for research. Thus, establishing new cell lines supports the 3R guidelines (24) and will help to achieve the future aim of ending the use of animals in research in the USA and the European Union (67). This new MC model will therefore sustain future studies on mechanisms and pharmacologic intervention of MC activation.

## Acknowledgements

This work was supported by grants from the Deutsche Forschungsgemeinschaft (DFG HU794/12-1, HU794/14-1, LI1045/6-1, WE2554/15-1, ME3431/2-1).

---

**Suppl. Fig. 1:**
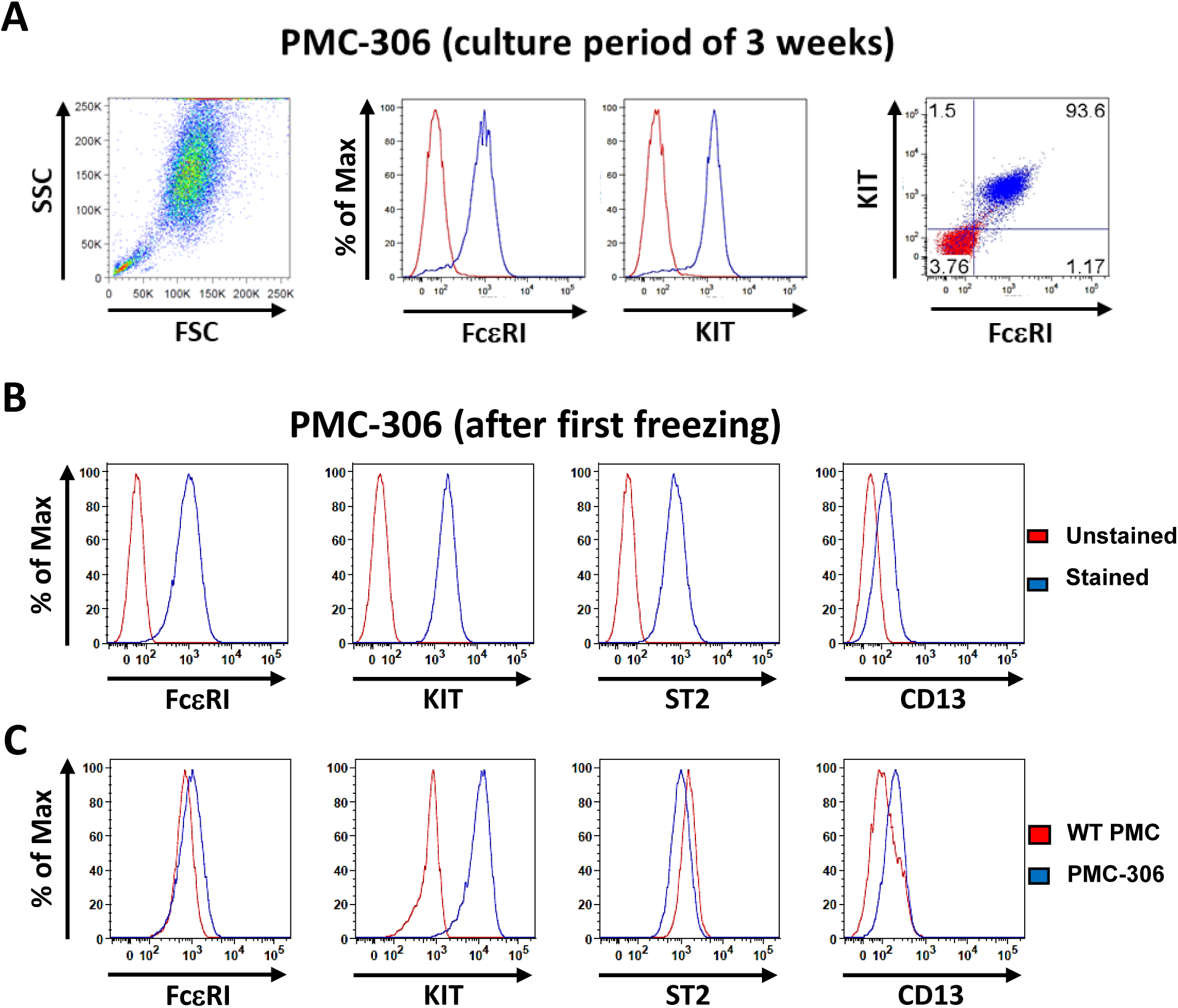
MC surface marker expression of PMC-306 before and after immortalization. **(A)** FACS analysis of PMC-306 cells before transformation after 3 weeks under regular PMC culture conditions. FSC/SSC dot plot shows a typical WT PMC population of 93.6% FcεRI and KIT double-positive cells. **(B)** Representative histograms of MC surface marker expression (FcεRI, KIT, ST2 and CD13) in PMC-306 analyzed by FACS after cryopreservation. **(C)** Representative histograms of MC surface markers expression analyzed by FACS comparing primary WT PMCs and PMC-306 cells.

**Suppl. Fig. 2:**
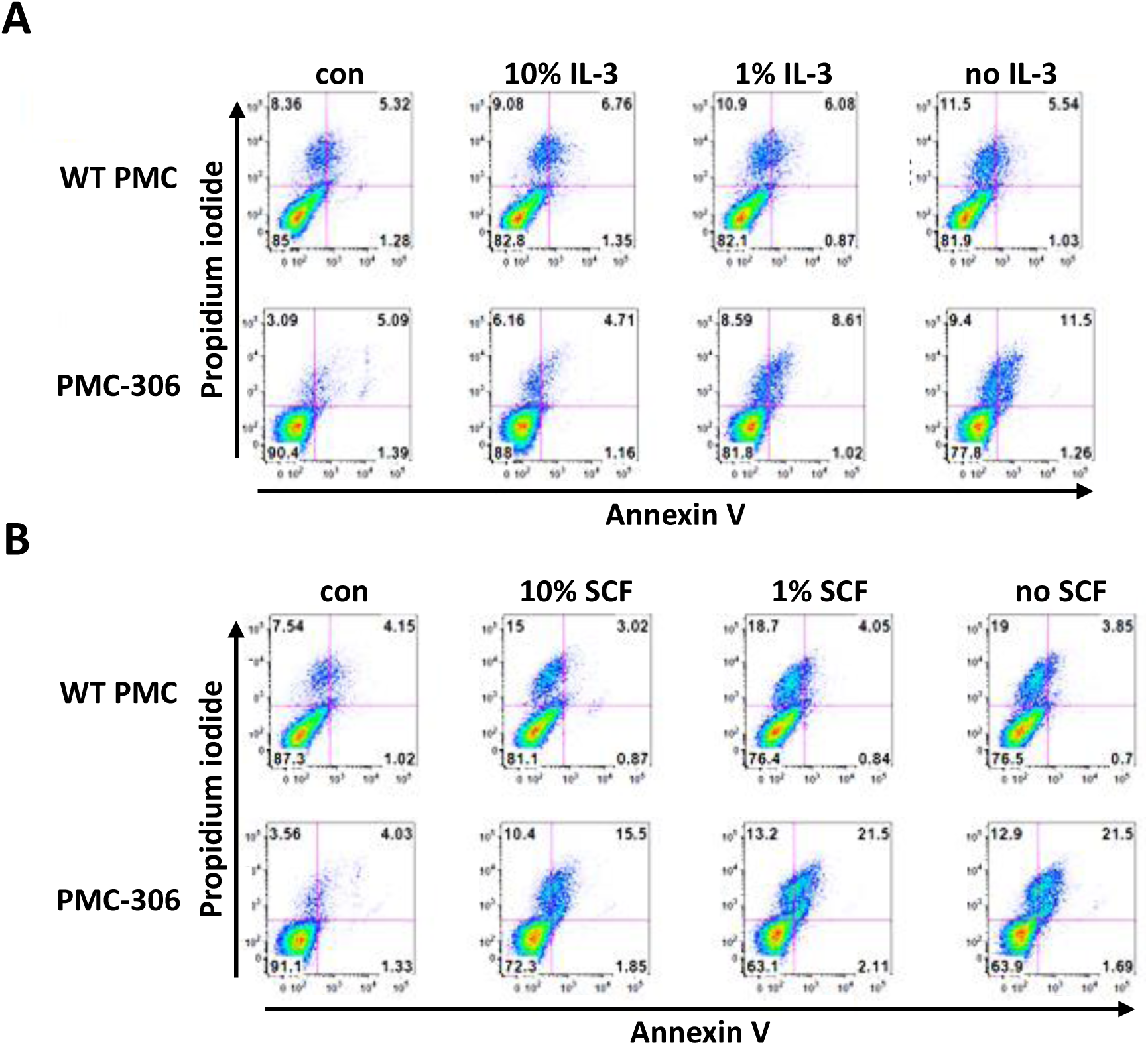
Analysis of primary WT PMC and PMC-306 viability under cytokine deprivation conditions. Representative FACS dot plots showing Annexin V and propidium iodide positivity in primary PMCs and PMC-306 cells under IL-3 **(A**) or SCF **(B)** deprivation for 72 hours. The percentage of single positive, double positive, and double negative cells is provided in the associated gates.

**Suppl. Fig. 3:**
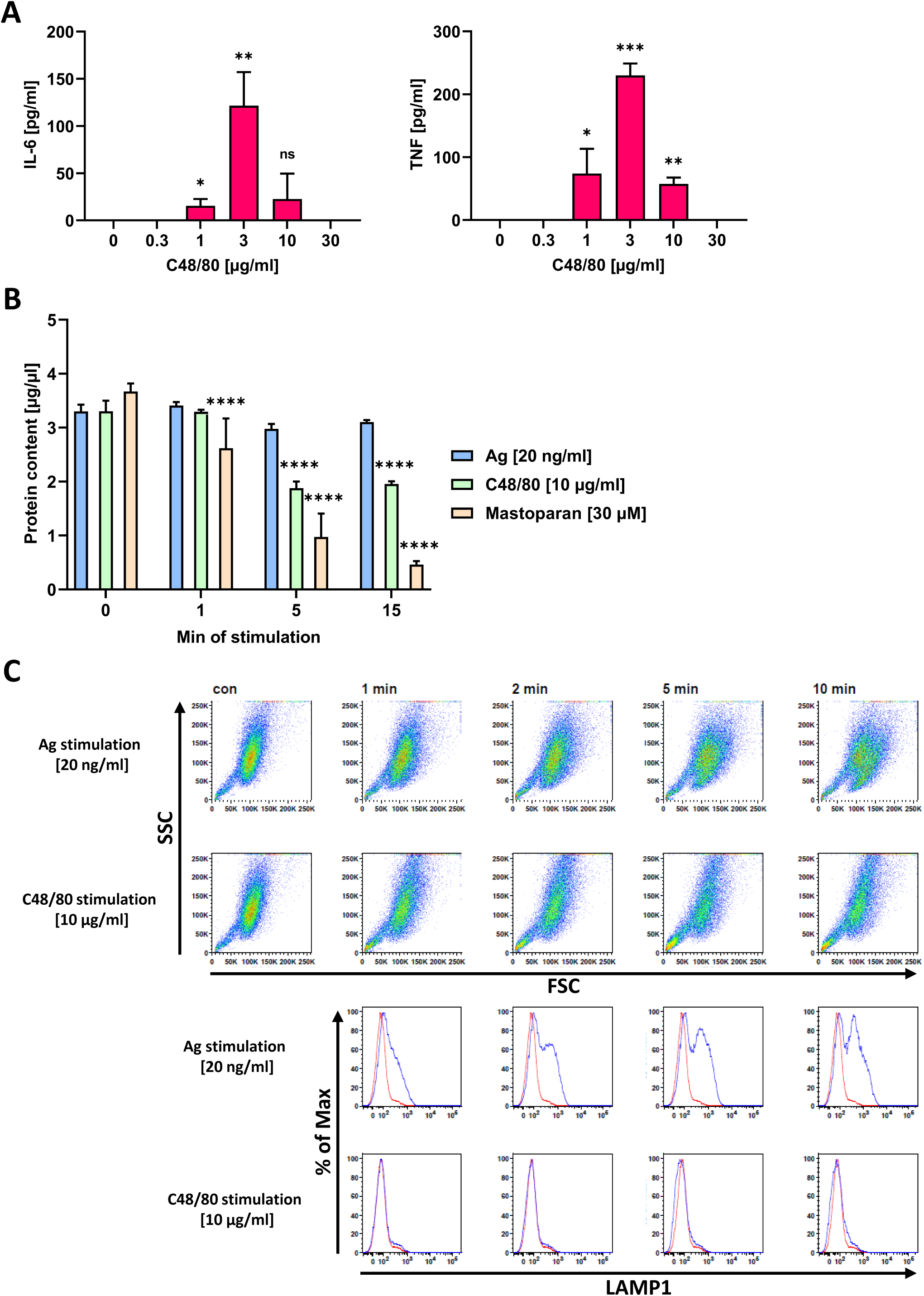
Analysis of MRGPRB2 activation shows unconventional degranulation and impact on cellular integrity. **(A)** ELISA measurement of secreted IL-6 (left) or TNF (right) from PMC-306 cells stimulated with increasing concentrations of C48/80 (n=4). **(B)** BCA assay to determine the protein concentration in cell lysates of PMC-306 cells stimulated with either Ag [20 ng/ml], C48/80 [10 µg/ml] or Mastoparan [30 µM] for the indicated time points (n=3). **(C)** Representative flow cytometry dot plots and histograms of primary WT PMCs stimulated with either Ag [20 ng/ml] or C48/80 [10 µg/ml] for indicated time points and stained with anti-LAMP-1 to determine granule externalization. Data are shown as mean +SD. **(A)** Ordinary one-way ANOVA followed by Dunnett multiple comparisons test. **(B)** Two-way ANOVA followed by Sídák multiple comparisons test. *p*>0.05 ns, * *p*<0.05, ** *p*<0.01, *** *p*<0.001, **** *p*<0.0001. Stars indicate the significance level relative to the respective controls.

**Suppl. Fig. 4:**
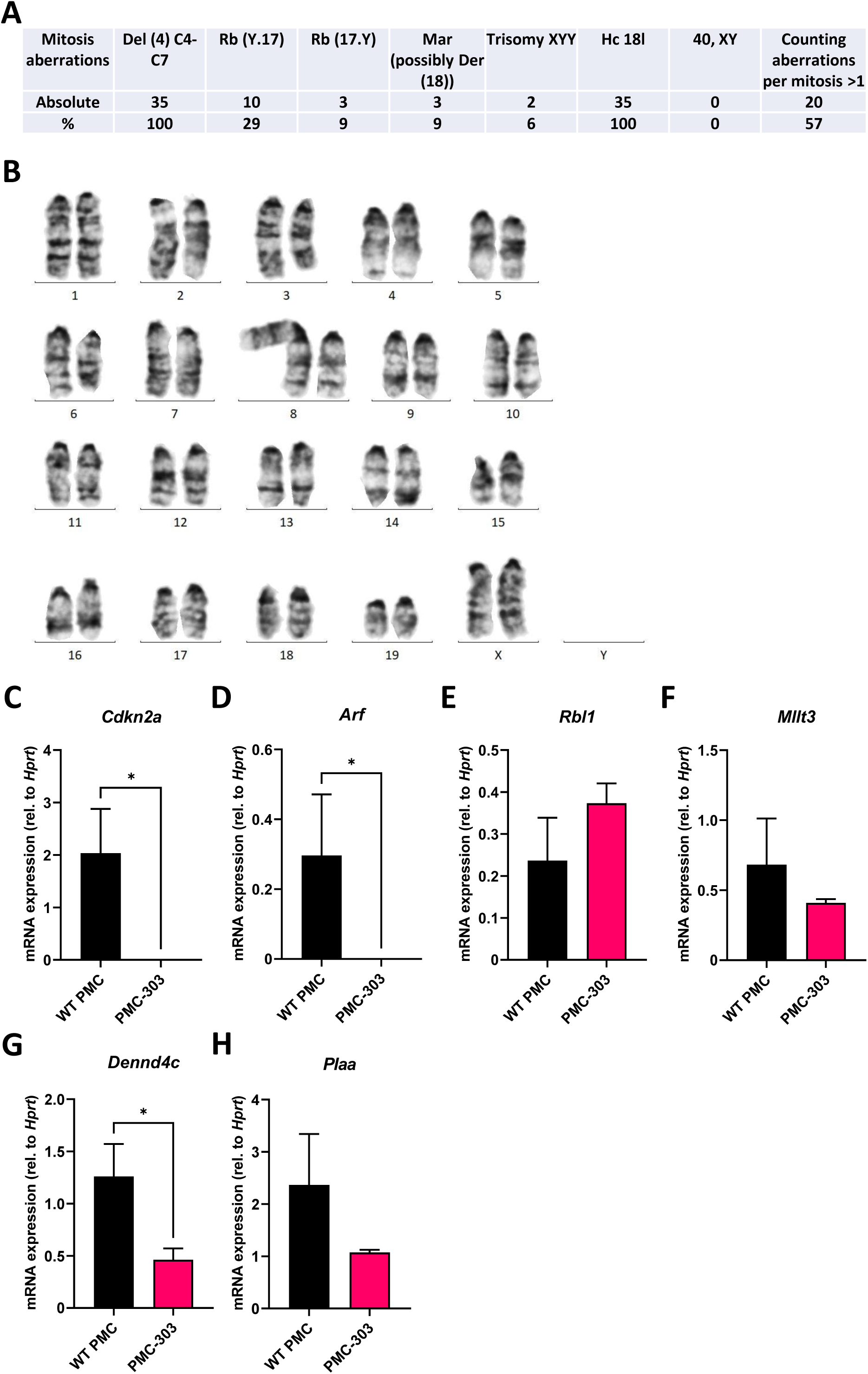
Cytogenetic and gene expression analysis of the independently transformed PMC cell line PMC-303. **(A)** Table of all discovered cytogenetic aberrations of mitotic chromosomes of the PMC-306 cell line with associated frequencies. **(B)** Representative G-banded karyotype of the PMC-303 cell line with trisomy 8 due to a homologous Robertsonian translocation of chromosome 8 (Rb(8.8)). **(D)**-**(H)** RT-qPCR analysis of genes encoded within in the region of chromosome 4, which is deleted in the PMC-306 cell line (Chr4 qC4-qC7). Gene expression of *Cdkn2a*, *Arf*, *Rbl1*, *Mllt3*, *Dennd4c* and *Plaa* were measured in PMC-303 cells in comparison to primary WT PMCs. Data are expressed as mean +SD. Unpaired, two-tailed Student’s *t*-test with Welsh’s correction. * *p*<0.05.

**Suppl. Fig. 5:**
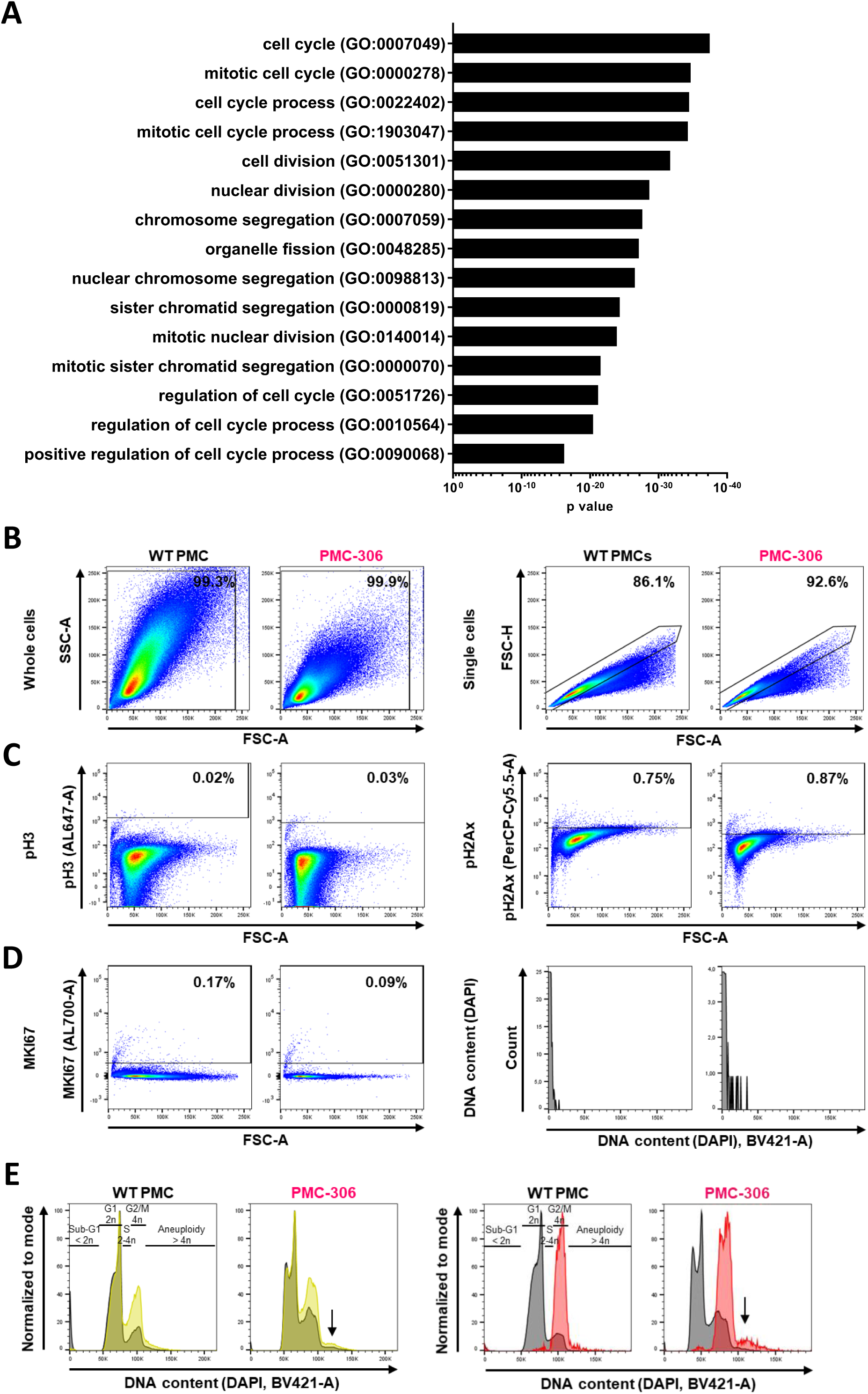
GO Term analysis of the NGS data set, gating strategy and background determination for flow cytometry (FACS) and analysis of aneuploidy. **(A)** For the GO term enrichment analysis all genes regulated at least 5-fold between primary WT PMCs and PMC-306 cells were included. The top 15 GO terms with the lowest p values are depicted. GO term analysis was performed using the gene ontology knowledgebase server (http://geneontology.org/) **(B-D)** Gating strategy and background determination for the transformed PMC-306 line (red) and primary WT PMCs (black). **(B)** Left panels: Forward *versus* side scatter (FSC *vs.* SSC) gating for identification of cells of interest. FSC indicates cell size, SSC indicates cell granularity. Right panels: FSC-A *vs.* FCS-H gating for identification of single cells. **(C,D)** Unstained controls, which were not incubated with antibodies to assess background for staining against phospho-histone 3 (AL647-A, pH3) and phospho-histone H2Ax (pH2Ax, PerCP-Cy5.5-A) **(C)**, MKI67 (AL700-A) and DAPI (DNA content, BV421-A) **(D)**. **(E)** Representative histograms of primary WT PMCs and PMC-306 cells stained with an antibody directed against MKI67 (AL700-A) and phospho-histone H3 (pH3, AL647-A). Cellular DNA content was determined by DAPI staining (BV421-A) for assignment to the distinct cell cycle phases as follows: SubG1: < 2n DNA content; G1: 2n; S: 2-4n; G2/M: 4n; Aneuploidy: > 4n. Representative overlay of FACS-histograms of DNA content in all single cells (black) *versus* MKI67+ (yellow, left) or pH3+ (red, right) single cells (normalized to mode). Black arrows indicate MKI67+ and pH3+ cells with a DNA content above 4n (aneuploidy).

## Notes

### Competing Interest Statement

The authors have declared no competing interest.

